# BCG vaccination reduces the rate of Mycobacterium tuberculosis dissemination between murine lungs

**DOI:** 10.64898/2026.03.10.710600

**Authors:** Dipanjan Chakraborty, Vitaly V. Ganusov

**Affiliations:** Host-Pathogen Interactions (HPI) program, Texas Biomedical Research Institute, San Antonio, TX 78227, USA; Molecular Microbiology and Immunology department, University of Texas at San Antonio, San Antonio, TX 78249

**Keywords:** Tuberculosis, Mycobacterium tuberculosis, mathematical model, within-host dynamics, BCG vaccine, disseminated disease

## Abstract

The BCG vaccine remains the only licensed vaccine against tuberculosis, yet the mechanisms underlying its protection are poorly understood. Using data from ultra-low dose (ULD) infection of control and BCG-vaccinated mice, we developed mathematical models of Mycobacterium tuberculosis (Mtb) growth and dissemination between the lungs. Alternative direct and indirect dissemination pathways fit unvaccinated data equally well, indicating multiple plausible routes of spread. Best fit models predicted that BCG reduces the rate of Mtb dissemination between lungs by nearly 90% while only modestly reducing the replication rate by 9%. Interestingly, stochastic simulations of Mtb dynamics with early bacterial killing showed that a growth-reducing vaccine would result in fewer infected mice, lower lung CFU, and decreased bilateral infection of the lung even at relatively moderate efficacy levels. Further, we estimated sample sizes needed to detect vaccine effects in preclinical trials. This framework provides a quantitative, mechanistic basis for interpreting BCG efficacy and offers a framework for evaluating next-generation TB vaccines in the ULD infection model.

## Introduction

Tuberculosis (**TB**), a disease caused by *Mycobactϵrium tubϵrculosis* (**Mtb**), remains one of the prevalent infectious diseases in humans^1^. Each year, approximately 10 million indivduals develop TB, and many individuals, particularly children and immunocompromised adults, develop extrapulmonary disease, in which Mtb disseminates beyond its primary site of infection, the lung^2,3^. Even though drug-susceptible TB can be effectively treated within 4-6 months in most individuals and relatively short and effective treatments are now available for patients with multi-drug resistant (**MDR**) TB^4^, controlling TB prevalence in humans will require additional tools such as an effective vaccine^5^.

The Bacillus Calmette–Guérin (**BCG**) vaccine, first used to immunize a human volunteer in 1921, remains the only licensed vaccine against TB in humans^6–8^. Initial randomized, placebo-controlled trials conducted in the UK and Scandinavia demonstrated substantial protection of BCG vaccine against pulmonary and extrapulmonary TB especially in children in 2-3 years of the follow up ^9,10^; however, later studies found more variable efficacy of BCG with some studies, such as the Chingleput study in India, reporting no protection against TB^11^. A recent meta-analysis of 18 clinical trials confirmed that BCG provides consistent and robust protection against severe, disseminated forms of childhood TB, including miliary TB and TB meningitis^12–15^. Yet, mechanisms by which BCG induces protection of children against TB remain poorly understood and typically are difficult to study in humans^16–18^.

Multiple animal species have been used to quantify BCG-mediated protection against Mtb infection and to identify mechanisms of such protection^18–20^. In general, intradermal BCG vaccination induces measurable protection of mice, rats, guinea pigs, rabbits, and macaques against Mtb challenge; vaccinated animals typically have reduced bacterial burdens in the lungs and/or extrapulmonary sites (e.g., spleen), reduced lung pathology, and longer survival^21–39^. In particular, in typical mouse strains such as C57BL/6 or BALB/c, BCG vaccination reduces lung bacterial loads by 0.5 − 1 log_10_, and guinea pigs (species highly susceptible to Mtb infection) after BCG vaccination have up to 2 log_10_ reduced lung CFU as compared to unvaccinated controls, measured several weeks after infection^21,31^. Interestingly, BCG vaccination can induce sterilizing protection in macaques, particularly following vaccination with high-dose intravenous inoculum^27^; protection against infection has also been reported with alternative vaccination routes such as pulmonary delivery of BCG^32^. Overall, previous studies have suggested that BCG is likely to mediate protection via a variety of different mechanisms, including generation of trained innate immunity, T cell memory, and antibodies^18,19,23,33,37,38^.

It has been well recognized that Mtb dose has a strong impact on the course of infection in animals (e.g., lung CFU numbers, lung pathology, and survival time)^28,40^; yet, whether degree of protection conferred by BCG vaccine depends on the Mtb exposure dose has remained relatively unexplored. Plumlee *ϵt al*. ^39^ found that exposing BCG-vaccinated B6 mice to a conventional dose (**CD**) of Mtb H37Rv (about 100 CFU/mouse) resulted in ∼1 log_10_ reduction in lung CFU as compared to unvaccinated controls at day 42 post infection; however, over time (at day 120) the difference in lung CFU between control and BCG-vaccinated mice became reduced and statistically non-significant. Interestingly, exposing control and BCG-vaccinated mice to an ultra-low dose (**ULD**) of Mtb H37Rv (with predicted effective dose of ∼ 1 CFU/mouse) resulted in a smaller proportion of mice with detectable CFU (i.e., infected), lower lung CFU (by about 1 log_10_ for 120 days), and less frequent bilateral infection of both right and left lungs^39^. However, this study did not investigate how BCG vaccination-impacted metrics such as infection rate, lung CFU, bilateral infection may be related; for example, whether reduction in the frequency of bilateral infection is simply a result of a lower lung CFU or if BCG reduces the per capita dissemination rate of bacteria between the lungs was not determined.

Here we used mathematical modeling to address these questions by studying impact of BCG vaccination on the probability of infection, Mtb growth in the lung and dissemination between right and left lungs in settings of ULD infection^39^. Interestingly, we found that mathematical models that assume direct (lung to lung) or indirect (lung to intermediate tissue to lung) dissemination pathways fit the data from unvaccinated animals with similar quality suggesting that existing data are insufficient to discriminate between the alternatives^41^. Yet, independently of the model of Mtb dissemination we found that BCG vaccination reduced both the rate of Mtb replication in the lung and the rate of Mtb dissemination between the lungs; alternative models in which BCG increased the Mtb clearance rate or had no impact on dissemination did not fit the data well. Importantly, the model predicted BCG-induced reduction in Mtb replication rate naturally leads to higher infection clearance probability (by 9%) and reduced (by about 1 log_10_) total CFU in the lung in vaccinated animals. We used our mathematical modeling-based framework to evaluate impact of growth-or migration-reducing vaccine on Mtb dynamics and dissemination and provided power analyses on the number of mice that would be required to detect efficacy of a hypothetical vaccine. Taken together, our work establishes a predictive, quantitative framework for interpreting the efficacy of the BCG vaccine and serves as a foundation for evaluating next-generation TB vaccines in ULD infection settings.

## Materials and Methods

### Experimental data

Plumlee *ϵt al*. ^42^ have developed a protocol in which a standard culture used for conventional dose (**CD**) infections is diluted ∼50-fold, filtered, and used in a nebulizer to produce ultra-low dose (**ULD**) aerosol to infect mice in the infection chamber. In a typical ULD experiment, about 60-80% of the mice get infected (as determined by CFU measured at later time points for all mice in the experiment), and the number of founding strains detected in individual mice using barcoded bacteria after ULD infection is consistent with a Poisson distribution^42^. In a recent study, Plumlee *ϵt al*. ^39^ used the same protocol to infect over 1,000 of B6 mice with an ULD of H37Rv Mtb (∼1 CFU/mouse, **Suppl. Fig. S1**A); approximately half of mice (*n* = 531) were subcutaneously immunized 8 weeks earlier with 10^6^ BCG-Pasteur and half were left unimmunized (*n* = 521). In total, the authors performed 21 experiments.

For most of our analyses, we included only infected mice (i.e., mice with CFU *>* 0 in **at least one of the** lungs); excluding uninfected mice or mice with no CFU data for right and left lungs yielded CFU data for *n* = 300 unvaccinated and *n* = 256 BCG-vaccinated mice (**Suppl. Fig. S2**). In the experiments, Plumlee *ϵt al*. ^39^ measured Mtb CFU at different days post-infection in the right (**RL**) and left (**LL**) lungs separately (**Suppl. Fig. S2**); our previous studies suggested that a bias in the initial infection towards the right lung most likely arises due to its larger volume/weight^42^ (**Suppl. Fig. S1**B). While infection could be initiated in any of the lungs (or in some cases in both lungs if there are 2+ bacteria entering the lung), it was difficult to envision one deterministic model that would incorporate such stochastic outcomes of the initial infection step. Therefore, to allow fitting ordinary differential equation(ODE)-based mathematical models to CFU data in the RL and LL we re-arranged the data by assigning larger CFU values (between RL and LL in individual mice) as Lung 1 (*L*_1_) and the smaller value as Lung 2 (*L*_2_), with the assumption that bacteria initiate the infection in the Lung 1 (with CFU = 1), and that the infection disseminates over time to from Lung 1 to Lung 2 (**Suppl. Figs. S2 and S3** and **Fig. 1**).

**Figure 1:**
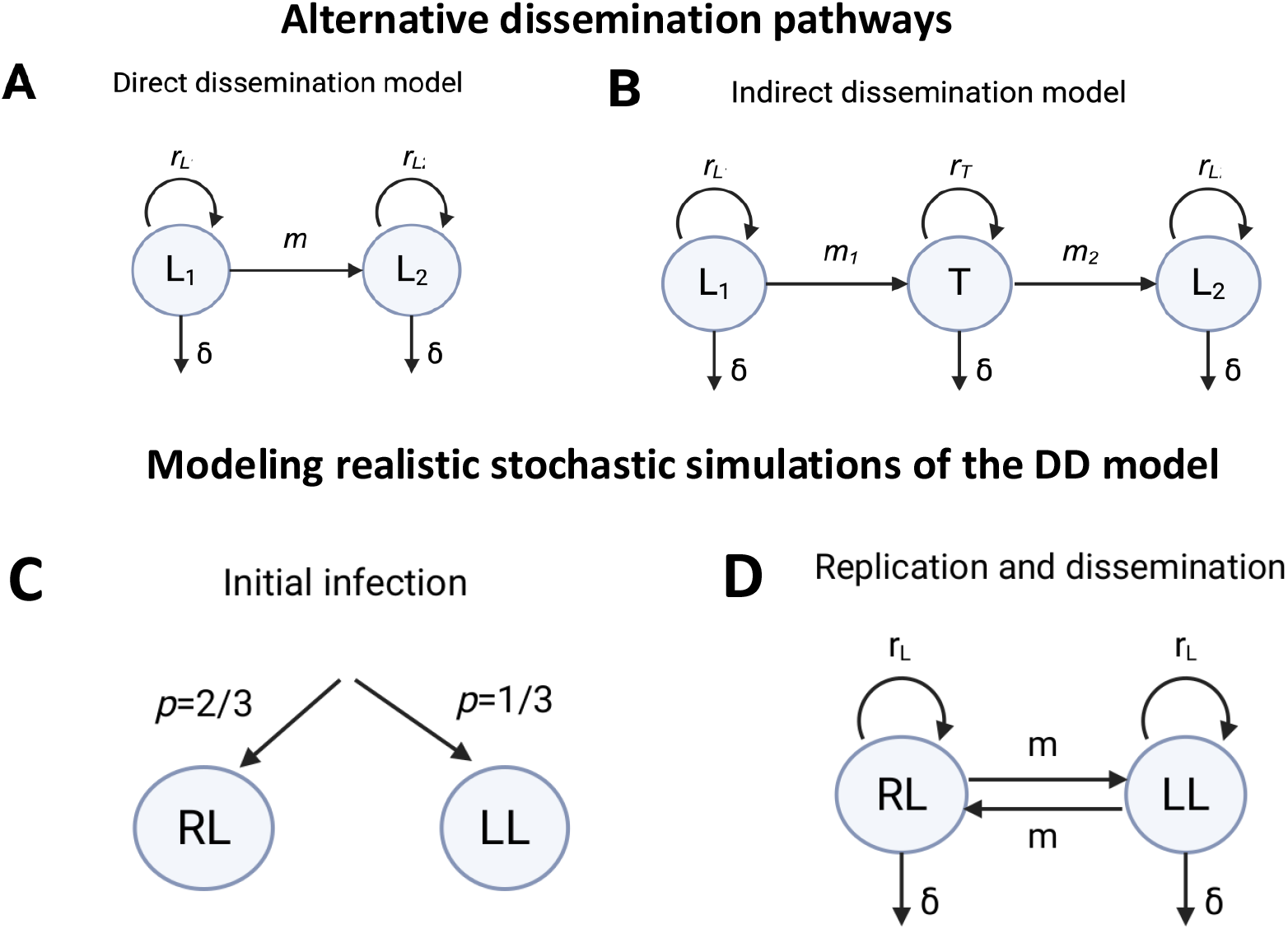
Schematics of alternative dissemination pathways of Mtb between the lungs of ULD-infected mice. In our main ODE model-based analyses we assume that infection with one bacterium (**Suppl. Fig. S1**A) starts in Lung 1 (*L*_1_) where bacteria replicate and die at rates *r*_*L*_ and *δ*, respectively. In the direct dissemination (**DD**) model (panel **A** and **eqns. (1)–(2)**) Mtb disseminates from Lung 1 to Lung 2 at a rate *m*, and in the indirect dissemination (**ID**) model (panel **B** and **eqns. (5)–(7)**), Mtb first disseminates from Lung 1 to an intermediate tissue *T* at a rate *m*_1_ and then from tissue *T* disseminates to Lung 2 at a rate *m*_2_. Mtb replicates and dies in both models in the Lung 2 at rates *r*_*L*_ and *δ*, respectively. In the ID model, Mtb replicates and dies in the intermediate tissue at rate *r*_*T*_ and *δ*, respectively. In our stochastic simulations (dubbed as “realistic” simulations, panels **C-D**) we allow the infection to occur with a Poisson-distributed number of bacteria (with average CFU = 1*/*mouse, **Suppl. Fig. S1**A) that infect the right lung (**RL**) or left lung (**LL**) with a probability 2/3 or 1/3, respectively (**Suppl. Fig. S1**B and **C**). Mtb then replicates and dies in the lungs at rates *r*_*L*_ and *δ*, respectively, and disseminates between the lungs at a rate *m* (**D**). In all models, we assume that average effective dose is 1 CFU.

### Mathematical models

Mathematical models, including those of within-host dynamics of bacterial infections, may include different levels of biological details. Which biological details to include in the model remains subjective but often relates to amount of quantitative data available. Because ULD infection of mice a novel infection model there is no comprehensive measurements of how different host cell populations behave during the infection; for example timing of recruitment of different circulating cells such as monocytes and neutrophils and later T cells, to the infection sites in the lung is unknown. A mathematical model that would explicitly include dynamics of such cell populations risks of overfitting the data. Therefore, here we adopted for an approach in which interactions between Mtb and host cells is reduced to time-dependent replication and death rates, with vaccination impacting these rates.

Specifically, we use a compartmental ODE-based modeling framework that focuses directly on predicting bacterial burden (CFU) over time^42–44^. Building on this approach, we explored several alternative mathematical models to describe Mtb dynamics in the initially infected lung (Lung 1) and its dissemination to the collateral lung, Lung 2 (**Fig. 1**A&B). We re-organized the data as Lung 1 and Lung 2 (see above) and assume that the infection is initiated in Lung 1 (with CFU = 1) but over time Mtb disseminates to Lung 2 (**Fig. 1** and see above). We consider two alternative pathways of Mtb dissemination between Lung 1 and Lung 2: (i) In the direct dissemination (**DD**) model (**Fig. 1**A), Mtb directly disseminates from the initially infected lung (Lung 1) to the collateral lung (Lung 2); (ii) in the indirect dissemination (**ID**) model (**Fig. 1**B), Mtb reaches the collateral lung (Lung 2) via an intermediate tissue (e.g., blood or spleen).

#### Direct dissemination (DD) model

We have used a simple system of linear ODEs to model the dynamics of Mtb (**Fig. 1**A&B):

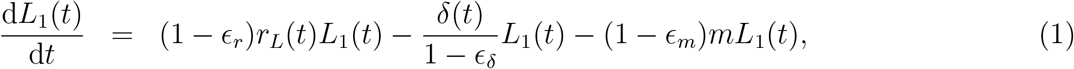

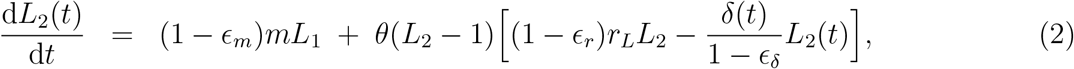

where *L*_1_(*t*) and *L*_2_(*t*) represent the Mtb numbers in Lung 1 and Lung 2 at time *t*; *r*_*L*_(*t*) denote the replication rate of Mtb; *δ*(*t*) is the death rate of Mtb; *m* is the migration rate of Mtb from Lung 1 to Lung 2 and *ϵ*_*r*_, *ϵ*_*m*_, and *ϵ*_*δ*_ quantify the effects of BCG vaccination on replication rate, migration rate, and death rate of Mtb, respectively. The Heaviside theta function *θ*(*x*) in **eqn. (2)** indicates that replication and death of Mtb begin in Lung 2 only when the number of disseminated Mtb exceeds 1 CFU. This modification was done to prevent rapid exponential growth of bacteria in Lung 2 when only a tiny fraction of a bacterium (≪ 1) is disseminated from Lung 1.

In this model formulation *ϵ*_*r*_ = 0, *ϵ*_*m*_ = 0, and *ϵ*_*δ*_ = 0 correspond to unvaccinated mice. For non-zero values of *ϵ*_*r*_, *ϵ*_*m*_, and *ϵ*_*δ*_, we can explore different models by choosing different combinations of *ϵ*_*r*_, *ϵ*_*m*_, *ϵ*_*δ*_. We note, however, that in addition of vaccination impact on Mtb dissemination rate, we consider vaccine impacting either only replication (*ϵ*_*r*_ *>* 0 and *ϵ*_*δ*_ = 0) or only death (*ϵ*_*r*_ = 0 and *ϵ*_*δ*_ *>* 0) but not both. The death rate, *δ*(*t*), is time-dependent but is kept the same for both Lung 1 and Lung 2.

To define how replication and death rates may change over the course of infection we adopted the approach outlined previously that assumed constant rates in a particular time period that could be different between different time periods^43,45^. In our preliminary analyses, however, we found that the time intervals provided previously (day 0-13, 13-26, 26-69, and 69-111) were not adequate for our data, for example, because our first measurement was for day 14 after infection, and we observed CFU decline between day 26 and 42 that was not captured in the longer previously defined time interval 26-69). After performing several regression analyses we oped for using time intervals slightly modified from the previous work: (day 0-16, 16-26, 26-42, and 42-125).

In our preliminary attempts of fitting models to data we found that we could not estimate both time-dependent Mtb replication and death rates together; therefore, we opted for fixing Mtb death rates to estimates provided previously^43^ with slight extension of the first time interval to include day 16:

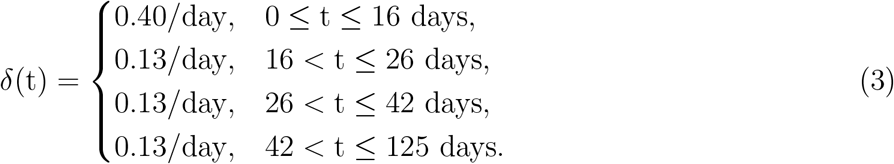

To maintain consistency with the time-dependent formulation of the death rates, we modeled the Mtb replication rates as time-dependent functions, denoted by *r*_*L*_(*t*):

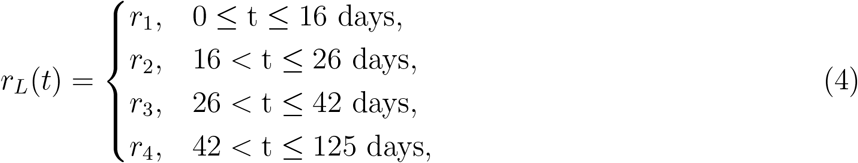

where we assume that the replication rate of Mtb at Lung 1 and 2 is the same. The migration rate of Mtb from Lung 1 to Lung 2 (*m*) is assumed to be independent of time. Each of the key terms in the model equations (**eqns. (1)–(2)**), the replication-driven growth term, the immune-mediated decay term, and the migration term representing dissemination is proportional to the bacterial load at a given time point, reflecting the assumption that these processes scale with the number of Mtb present. Since this DD model mimics the ULD Mtb infection in mice, we started our simulation with 1 bacterium (as the average effective dose is 1 CFU, **Suppl. Fig. S1**) in Lung 1 and 0 bacterium in Lung 2.

#### Indirect dissemination (ID) model

In the indirect dissemination model, the spread of Mtb to the collateral lung occurs through an intermediate tissue, e.g., blood or spleen (**Fig. 1**B). Therefore, the ID model has another equation for the Mtb dynamics in the intermediate compartment. If the intermediate tissue is the spleen, that allows for Mtb replication, the equations describing Mtb dynamics in Lung 1, tissue, and Lung 2 will be the following:

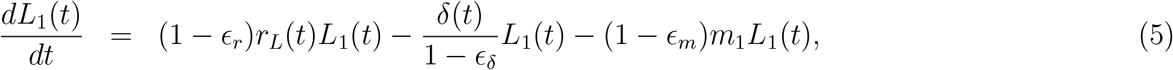

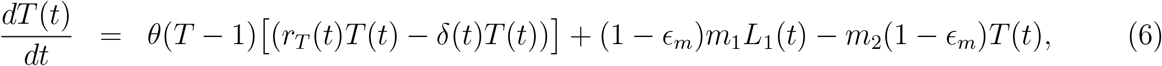

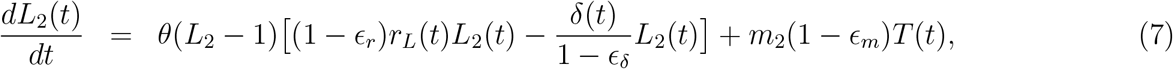

where *r*_*T*_ (*t*) indicates the Mtb replication rate at the intermediate tissue T, *m*_1_ is the migration rate of Mtb from Lung 1 to tissue, and *m*_2_ is the migration rate of Mtb from tissue to Lung 2. If the tissue is blood, then there will be no replication and death of Mtb within the blood, i.e., *r*_*T*_ = *δ*_*T*_ = 0; only the migration term would exist, signifying blood as a carrier of Mtb but not the place of extensive replication (**eqns. (S.1)–(S.3)**). The initial condition for starting the simulation is kept the same as described in the DD model. Further, the replication and death of Mtb in the tissue and Lung 2 would only occur when the number of disseminated Mtb crosses 1 CFU, as indicated by the Heaviside function *θ*(*x*) in **eqns. (6)–(7)**.

#### Fitting the direct dissemination (DD) and indirect dissemination (ID) models to data

To fit both the DD and ID models to the CFU data in mouse lungs we used nonlinear least-squares minimization with the **Levenberg–Marquardt (Marq)** optimization algorithm implemented in the `modFit` routine of the `FME` package in `R`. We log-transformed model predictions and the data and minimized the sum of squared residuals to find the best fit. Prior to analysis, mice with no detectable Mtb burden in whole lung were removed from the dataset. Consequently, a CFU value of zero occurred only when one lung was infected, and the contralateral lung had not yet acquired bacteria. Because these zero measurements cannot be log-transformed, all data were shifted by adding one CFU, such that

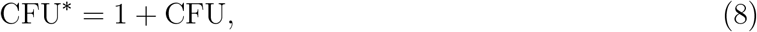

so that an experimental observation such as (*L*_1_ = 5, *L*_2_ = 0) becomes (*L*_1_ = 6, *L*_2_ = 1) prior to log transformation. This shift ensures that log_10_(CFU^∗^) remains well-defined while maintaining the biological ordering of the observed infection levels.

To initialize the ODE solver in R consistently with the experimental data, a data point at *t* = 0 was introduced which is the initial condition that we assumed but the associated 0 residual was later removed in calculating various additional characteristics (e.g. AIC) ^46^. To compare the DD and ID models, we used the Akaike Information Criterion (**AIC**) and the associated Akaike weights (*w*)^47^. The model with the lowest AIC (and thus highest Akaike weight) was selected as the most plausible among alternative models under consideration.

#### Stochastic simulations of the *L*_1_ → *L*_2_ version of the DD model

To model Mtb dynamics stochastically we converted the ODE version of a model (e.g., DD model) into discrete probabilistic events where Mtb replication, death, and migration occur with time-dependent transition rates derived from the best-fitted parameters of the ODE model (**Tab. 1**). These transition rules (**Tab. 1**) define how the system evolves through random events rather than via continuous population changes. For each simulation, the system was initialized with a single bacterium in Lung 1 and none in Lung 2 (**Suppl. Fig. S1**A). Because stochastic realizations yield only one possible outcome of a random process, multiple independent trajectories (typically 100-1,000) were generated to assess whether the variability captured by the model matches the experimental CFU distributions across lungs. Simulations were performed for 125 days using Gillespie’s exact stochastic simulation algorithm, implemented in the `GillespieSSA2` package in R^48,49^ and recorded with a step size of Δ*t* = 1.0 day.

**Table 1:** List of reactions and their rates of the stochastic simulations of the *L*_1_ → *L*_2_ version of the DD model. In the model, increase or decrease in the number of bacteria in Lung 1 occurs at the rate *r*_*L*_*L*_1_ or *δL*_1_, respectively, and migration of bacteria from Lung 1 to Lung 2 occurs at the rate *mL*_1_ (**eqns. (1)–(2)**). Note that the rates of Mtb replication and death in the Lung 1 and 2 are time-dependent (**eqns. (3)–(4)**).

| Reaction | Rate of Reaction |
| --- | --- |
| $L_1 \rightarrow L_1 + 1$ | $r_L L_1$ |
| $L_1 \rightarrow L_1 - 1$ | $\delta L_1$ |
| $L_1 \rightarrow L_1 - 1, L_2 \rightarrow L_2 + 1,$ | $m L_1$ |
| $L_2 \rightarrow L_2 + 1$ | $r_L L_2$ |
| $L_2 \rightarrow L_2 - 1$ | $\delta L_2$ |

#### Realistic stochastic simulations of the RL += LL version of the DD model

In the *L*_1_ → *L*_2_ version of the DD model we assumed a unidirectional dissemination pattern where bacterial infection was initiated in Lung 1, and it subsequently spread to Lung 2 (**Tab. 1**). However, in reality, the primary site of infection may arise in either left or right lung, followed by dissemination to the contralateral lung (**Fig. 1C&D**), and the initial number of inhaled bacteria could be higher than 1 CFU due to random nature of infection (**Suppl. Fig. S1**A). To incorporate these additional biological details, we modified stochastic simulations of the DD model to allow for randomness of lung infection and bidirectional migration between the two lungs (**Tab. 2**).

**Table 2:**
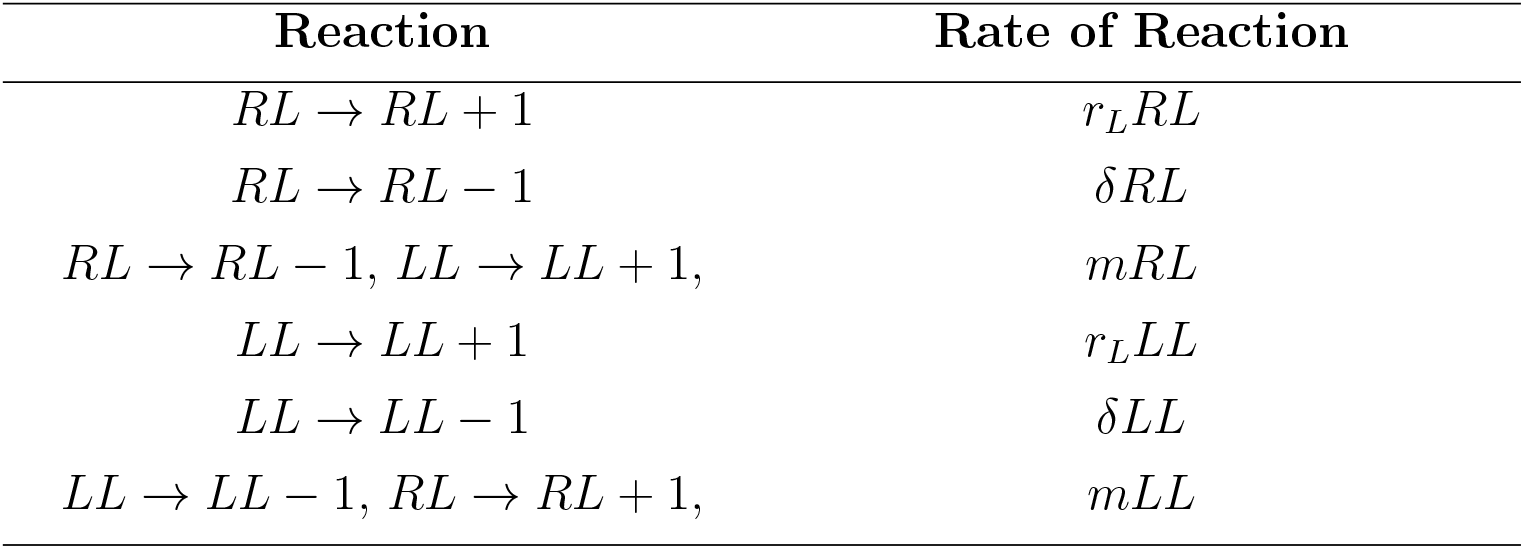
List of reactions and their rates of the realistic stochastic version of the DD model. In this model, Mtb infection is initiated in the right lung (RL) or the left lung (LL) with a probability 2*/*3 : 1*/*3 (**Fig. 1**C). Thereafter, the increase or decrease in the number of bacteria in RL occurs at a rate *r*_*L*_*RL* or *δRL*, respectively. Migration of bacteria from RL to LL and LL to RL occurs at a rate *m*.*RL* or *mLL* respectively. Replication rates and death rates of Mtb in the RL and LL are assumed to be the same as those of Lung 1 and Lung 2 (**eqns. (3)–(4)**).

First, we modeled the initial bacterial inoculum as a random draw from a Poisson distribution with mean *λ* = 1 (or any other dose), consistent with experimental measurements of infection dose variability (**Suppl. Fig. S1**A). Second, based on previous experimental observations^42^, we introduced an asymmetry in the initial infection probability, assigning a 2/3 chance that a bacterium lands in the right lung and a 1/3 chance for it to land in left lung (**Fig. 1**C and **Suppl. Fig. S1**B). Finally, we used parameters for replication (*r*_*L*_), death (*δ*), and migration (*m*) as estimated from fitting ODE version of the DD model to data to simulate Mtb dynamics stochastically in a similar fashion as was described above (e.g., **Fig. 1**C&D).

#### Quantifying efficacy of hypothetical vaccines in restricting bacterial dissemination

Using realistic stochastic simulations of the DD model (**Fig. 1**C&D and **Tab. 2**) we evaluated how a hypothetical vaccine influences the spread of Mtb between the lungs. We quantified the dissemination outcomes under two vaccine efficacy parameters: *ϵ*_*r*_, which reduces the rate of bacterial replication, and *ϵ*_*m*_, which reduces the rate of inter-lung dissemination. We assessed the probability that a mouse develops a bilateral infection, a clinically relevant marker of dissemination severity. We initialized simulations with a virtual cohort of 100 mice, using parameter values derived from the best-fitted model. For each parameter setting (e.g., varying *ϵ*_*r*_ while fixing *ϵ*_*m*_), we calculated the fraction of mice that developed bilateral infections at day 14 post infection relative to the total number of infected mice (with limit of detection LOD = 1). Because stochastic models produce variability across realizations, we performed 100 independent simulations for each combination of *ϵ*_*r*_ and *ϵ*_*m*_. The results were then averaged to obtain estimate of bilateral infection probability in different levels of vaccine efficacy.

#### Quantitative assessment of stochastic model fit using the Kolmogorov-Smirnov (KS) test

To evaluate the degree of agreement between stochastic simulation trajectories and experimental data, we propose to use the Kolmogorov–Smirnov (**KS**) test and KS statistics^50^. This non-parametric test quantifies the maximum distance between the cumulative distribution functions (**CDFs**) of the simulated and experimental data. In this context, the KS statistic (denoted as *D* ∈ (0, 1)) serves as a measure of the goodness of fit: smaller values indicate closer correspondence between model predictions and observed outcomes. We calculated an average KS distance 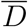 across multiple measurement days to account for temporal variability in bacterial counts as follows

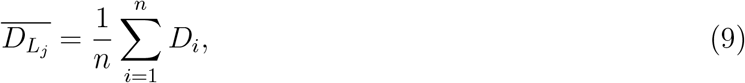

where *i* indexes the measurement days, *n* is the total number of days, and *j* denotes the lung under consideration (e.g., *j* = 1, 2 for Lung 1 and Lung 2 or RL and LL). A lower value of 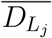 implies improved model fidelity. In the idealized case, 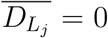 would indicate perfect agreement; however, due to intrinsic stochasticity and biological variability, such an outcome is not expected.

#### Model-based power analysis for detecting vaccine effects

Statistical power analysis determines the minimum number of animals needed to detect a treatment effect with a desired confidence. Plumlee *ϵt al*. ^39^ estimated the sample sizes required to distinguish impact of a vaccine on probability of infection at different levels of statistical power. Building on this idea, we performed a model-based power analysis using our realistic stochastic simulations of the DD model to assess how vaccine efficacy influences these sample size requirements. We examined two types of vaccines: vaccine reducing Mtb replication rate (*ϵ*_*r*_) or vaccine reducing Mtb dissemination rate (*ϵ*_*m*_). In the first case, we computed the infection probabilities in unvaccinated and vaccinated groups of mice by varying the efficacy of vaccination in replication (*ϵ*_*r*_). For a given sample size, these probabilities were summarized using a 2 *×* 2 contingency table, and statistical significance of group differences was evaluated using the *χ*^2^ test (*p* ≤ 0.05). By iterating across sample sizes, we identified the minimum cohort size required to achieve 80% power. Similarly, to assess dissemination control, we performed power analysis by examining the frequencies of bilateral lung infections in unvaccinated and vaccinated mice, under different levels of vaccine efficacy in blocking dissemination rate (*ϵ*_*m*_). The outcomes were organized into contingency tables, and *χ*^2^ testing was again applied to assess statistical significance across unvaccinated and vaccinated groups. In the calculation of power analysis, while exploring the effect of *ϵ*_*r*_, we set *ϵ*_*m*_ = 0, and similarly, when varying *ϵ*_*m*_, we fixed *ϵ*_*r*_ = 0.

## Results

### Mathematical modeling suggests multiple plausible routes of Mtb dissemination

Plumlee *ϵt al*. ^39^ exposed control or BCG-vaccinated B6 mice to an ultra-low dose (**ULD**) of Mtb and measured CFU in the right and left lungs at different times after infection; in total of 21 experiments, over 1,000 mice were used. Early after the exposure, bacteria were found primarily in one of the lung but over time, both lungs became infected. It is well understood that in mice, Mtb typically disseminates to extrapulmonary sites (e.g., lymph nodes or spleen) ^51–53^; however, whether systemic dissemination is required for infection of both lungs especially in settings of ULD infection has not been determined.

Mtb dynamics at ULD infections is intrinsically stochastic due to a low number of founding bacteria^42^ and we initially could not come up with a simple deterministic (ODE-based) mathematical model to describe Mtb dynamics and dissemination between the right and left lungs. Because in Plumlee *ϵt al*. ^39^ experiments, the average effective dose was 1 CFU/mouse (**Suppl. Fig. S1**A), per Poisson distribution most infections (∼60%) would be initiated by a single bacterium landing in the right or left lung. Therefore, we opted for a simplified mathematical model that assumes that infection is initiated by a single bacterium in one of the lungs (denoted as Lung 1) and over time, the bacteria disseminate to the collateral Lung 2. The dissemination may be direct or indirect (**Fig. 1**A-B): the direct dissemination (**DD**) model assumes Mtb migration directly from Lung 1 to Lung 2, while the indirect dissemination (**ID**) model assumes that Mtb passes via an intermediate tissue; Mtb may be replicating in the intermediate tissue (e.g., spleen) or just being transported between the lungs (e.g., via blood).

To fit the models to data we re-classified CFU in the right or left lung as CFU in Lung 1 or 2 depending on the CFU value in the two lungs for each individual mouse (i.e., larger CFU value was assigned to Lung 1 and the smaller value to Lung 2, **Suppl. Figs. S2 and S3**). This procedure is likely accurate when CFU in one of the lungs is much larger than in another but may introduce noise when CFU numbers in both lungs are similar. Thereafter, we fit the DD model (**eqns. (1)–(2)**) to the modified data on CFU in Lung 1 and Lung 2 (see Materials and methods for details); the model accurately captured rapid rise in CFU in Lung 1 followed by bacterial control and decline in CFU after the peak, and then slow increase in CFU over time (**Fig. 2**A). The model also accurately predicted continuous accumulation of CFU in the collateral lung (Lung 2) over time (**Fig. 2**B). In the model, CFU dynamics was determined by changes in the Mtb replication rate that had relatively high values early in the infection (*r*_*L*_ ≈ 0.94*/*day), declined to low levels (*r*_*L*_ ≈ 0.1*/*day), but then increased in the chronic phase (after 40 days *r*_*L*_ ≈ 0.14*/*day) remaining slightly higher than the death rate leading to slow increase in CFU number over time (**Fig. 2**C & **Suppl. Tab. S1**). Estimated Mtb per capita dissemination rate *m* = 2.7 *×* 10^−3^*/*day appears to be quite small yet sufficient to cause Mtb dissemination to the collateral lung by 2-3 weeks post infection (**Fig. 2**B). Interestingly, allowing for the Mtb replication rate to differ between Lung 1 and Lung 2 did not improve the model fit (F-test for nested models, *p* = 0.24), suggesting that changes in the rate of Mtb replication with time in both lungs may be set at the time of infection.

**Figure 2:**
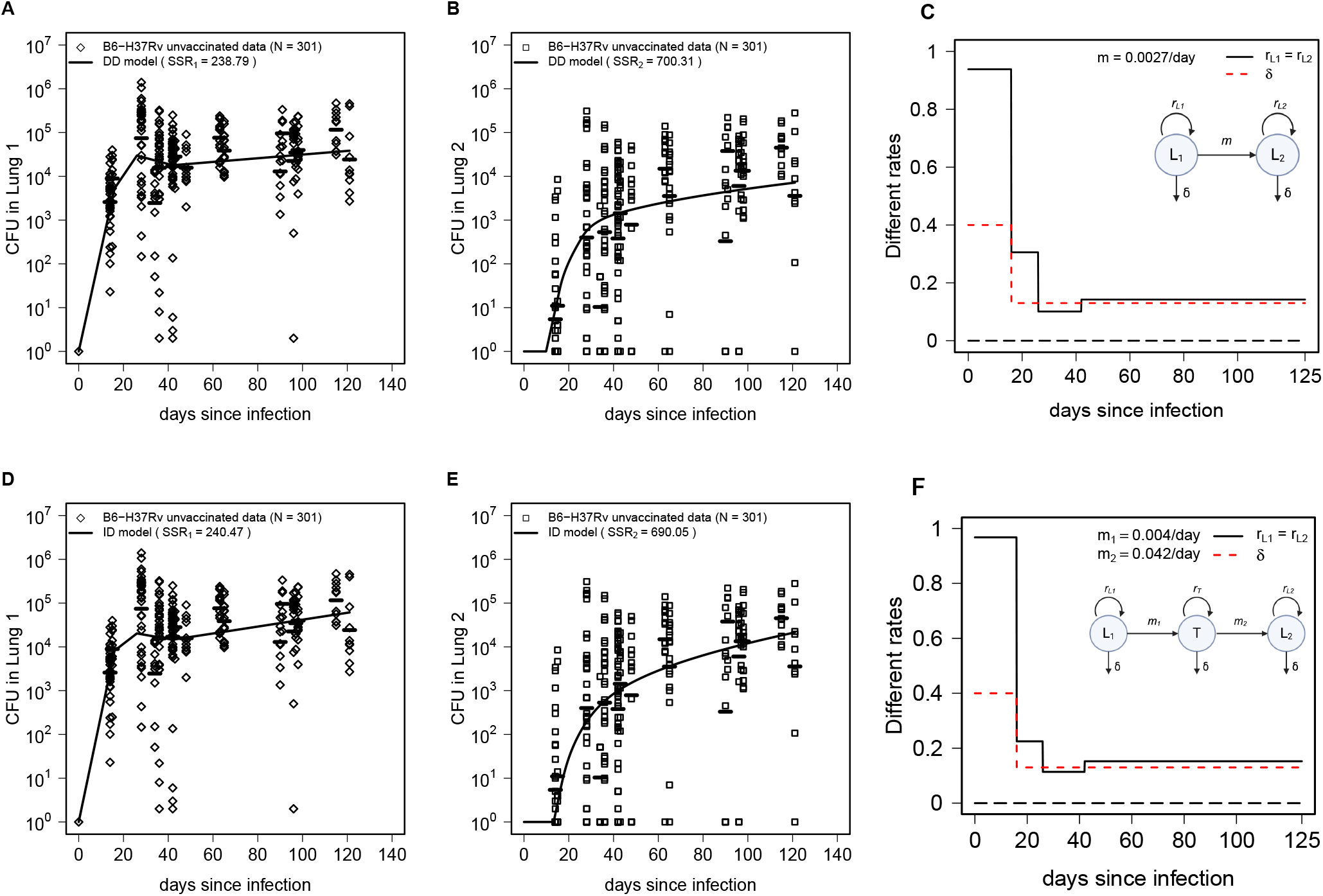
Alternative mathematical models assuming direct or indirect Mtb dissemination between murine lungs describe the data on Mtb dynamics in unvaccinated mice with similar quality. We fit the direct dissemination model (**eqns. (1)–(2)**, **Fig. 1**A, **A-C**) or indirect dissemination model (**eqns. (5)–(7)**, **Fig. 1**B, **D-F**) to CFUs found in lungs of unvaccinated mice (after re-arranging the data as Lung 1 and Lung 2, see Materials and methods for detail). We show the data as markers and model fits as lines in A-B and D-E along with estimated rates of Mtb replication in the lung (C&F) or in the intermediate tissue (**Suppl. Figs. S4 and S5** and **Suppl. Tab. S1**). In the model fits we assumed *L*_1_(0) = 1 and *L*_2_(0) = 0. The quality of the model fit was evaluated by sum of squared residuals (**SSR**) or AIC (see **Suppl. Tab. S2**). The total SSR is 939.1 and 930.52 for DD and ID models, respectively, and AIC values are 277.8 and 282.5 for DD and ID models, respectively.

We next fit the ID model (**eqns. (5)–(7)**) that assumes that Mtb is able to replicate in an intermediate tissue (**Fig. 1**B), to the data. Interestingly, the model could also accurately describe CFU in the Lung 1 and 2 at somewhat similar rates of Mtb replication (**Fig. 2**D-F & **Suppl. Tab. S1**); the model predicted relatively slow migration rate from Lung 1 to the intermediate tissue but then rapid migration from the tissue to Lung 2 (**Fig. 2**F). The model fits also predicted relatively rapid Mtb replication in the intermediate tissue (**Suppl. Fig. S4**A); interestingly, the model-predicted CFU in the tissue was consistent (but slightly lower than the mean) with the independently measured bacterial numbers in the spleens of ULD-infected mice at day 63 post infection (**Suppl. Fig. S4**B). Importantly, an alternative version of the ID model that assumes that dissemination between murine lungs occurs by passive transport (without replication), e.g., via blood, also fit the data well (**Suppl. Fig. S5**). Thus, our analysis suggests that both direct and indirect dissemination routes of Mtb spread between individual lungs are possible, and additional data would be required to discriminate between alternatives^41^. Statistically, however, the DD model fit the data with slightly better quality (based on AIC and Akaike weights) in part because of the smaller number of model parameters (**Suppl. Tab. S2**).

### BCG vaccination reduces Mtb replication and dissemination rates

We next turned to evaluating impact of BCG vaccination on Mtb dynamics and dissemination in ULD-infected mice. Visual inspection of the data indicated lower CFU in both right and left lung (or Lung 1 and 2, **Suppl. Figs. S2 and S3**) suggesting impact on net Mtb growth rate. To minimize the number of parameters fitted to data (and thus to reduced the likelihood of overfitting) we opted for the approach in which we fixed estimates of Mtb replication and dissemination rates to values estimated from unvaccinated mice but allowed the overall vaccine efficacy at reducing Mtb replication rate *ϵ*_*r*_ or dissemination rate *ϵ*_*m*_ (see **eqns. (1)–(2)**), and fitted the model by varying only *ϵ*_*r*_ and/or *ϵ*_*m*_. Interestingly, we found that in order to accurately fit the data, both DD and ID models required that BCG vaccination reduces both Mtb replication and dissemination rates (**Fig. 3**): the DD model predicted 9% and 89% reduction in the Mtb replication and dissemination rates, respectively, while the ID model (with lymph node/spleen being intermediate tissue) predicted rate of 9% and 65% reduction in replication and dissemination rate, respectively (**Fig. 3**). In the ID model Mtb disseminates via two sequential steps (Lung 1-tissue-Lung 2), each attenuated by the same factor (1 − *ϵ*_*m*_); thus, the net probability of Mtb dissemination is reduced by (1 − (1 − *ϵ*_*m*_)^2^) ≈ 88%, closely matching the 89% reduction estimated directly by the DD model. The ID model more accurately fit the data on Lung 1, while the DD model better fit the data for Lung 2 (note SSR values in **Fig. 3**). Statistically, the DD model fit the data slightly better (based on AIC values, **Suppl. Tab. S2**).

**Figure 3:**
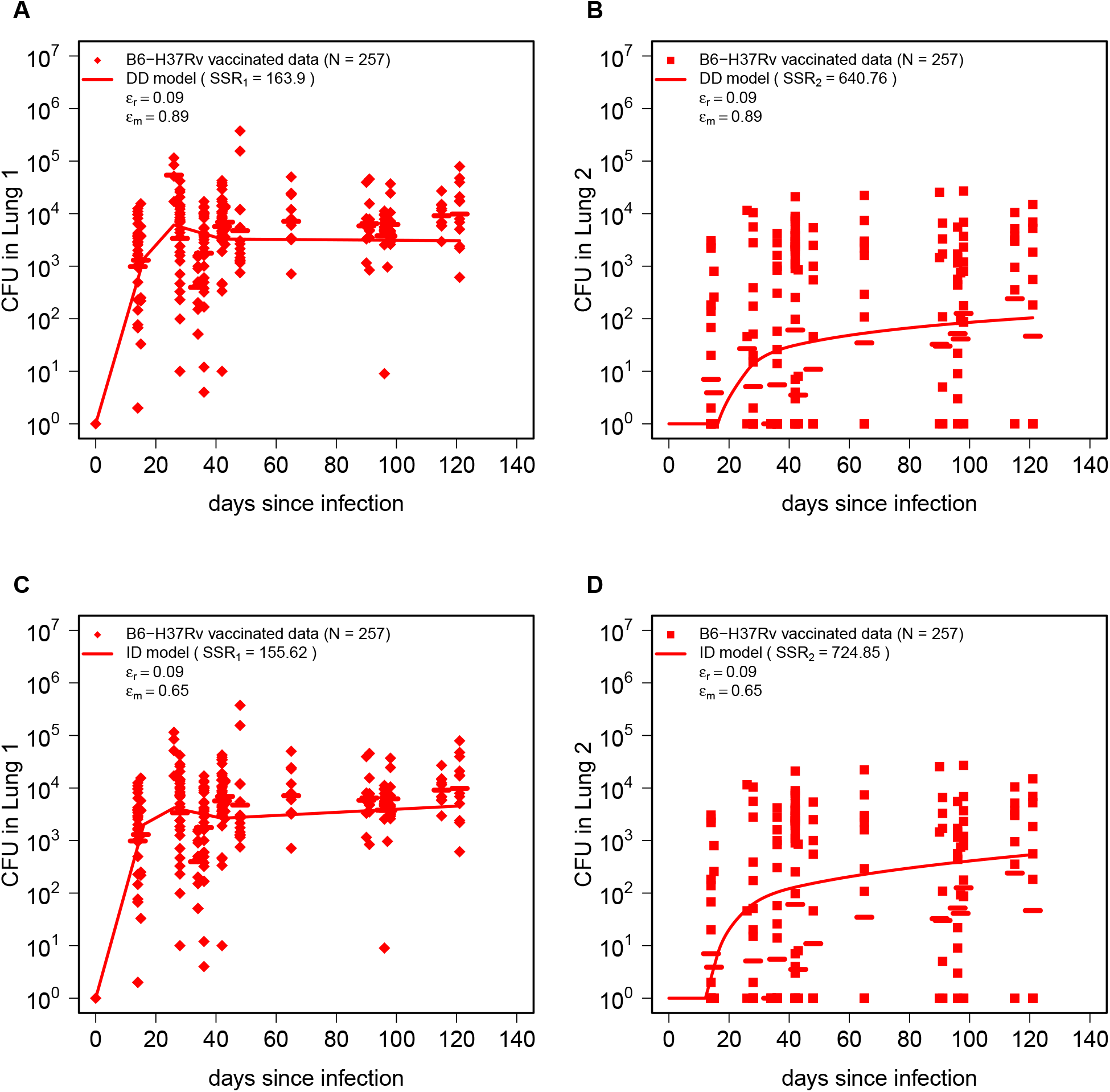
Both direct and indirect dissemination models predict that BCG vaccination significantly reduces Mtb dissemination rate and slightly reduces Mtb replication rate. Using the estimates of Mtb replication (*r*_*L*_) and dissemination (*m*) rates for unvaccinated mice (**Fig. 2**) we fit the DD (**eqns. (1)–(2), A-B**) or ID (**eqns. (5)–(6), C-D**) models to CFU data on Mtb dynamics in Lung 1 (**A&C**) and Lung 2 (**B&D**) in BCG-vaccinated mice. The data are shown by markers, and best-fit model predictions are shown by lines. The estimated efficacy of BCG vaccine at reducing the rate of Mtb replication (*e*_*r*_) and Mtb dissemination rate (*e*_*m*_) is shown on individual panels (see also **Suppl. Tab. S2**).

We noted that while the DD model accurately predicted average CFU in Lung 1 in the first 45 days post infection, the model consistently predicted lower average lung CFU than that observed experimentally (**Fig. 3**A&C). We wondered if this could arise if efficacy of BCG vaccine at reducing the rate of Mtb replication may be time-dependent. We therefore re-fitted the DD model by allowing *ϵ*_*r*_ to change between different time intervals (i.e., see **eqn. (3)**). Surprisingly, the model fits predicted decline of BCG vaccine efficacy with time since infection being 8-15% in the first 4 weeks of infection but then declining to 0.2-3% at later times (**Suppl. Fig. S7** A & B). Thus, ULD infection of mice also recapitulates decline in efficacy of BCG vaccine with time seen in humans^54^; however, the time scale of this decline is not comparable (months in mice vs. years in humans).

Our results show that, independently of the assumed mode of Mtb dissemination, BCG vaccination reduces the Mtb replication rate in the lung; this finding is important for two reasons. First, it naturally explains lower CFU numbers observed in BCG-vaccinated mice. While the average 9% reduction in replication rate may appear small, in about 25 days such a difference in replication rates will result in 10 fold difference in CFU numbers as observed experimentally (**Suppl. Fig. S2**). Second, *ϵ*_*r*_ = 9% reduction in the rate of Mtb replication naturally results in an identical increase in the probability of infection clearance when exposed to a dose of 1 CFU (infection extinction probability for 1 CFU is *p*_0_ = *δ/r*); thus, our analysis of the data from only infected mice made a similar inference of reduced infection probability of BCG-vaccinated mice as was done by Plumlee *ϵt al*. ^39^ by looking at the proportion of infected mice in the whole dataset.

Along with the models that fit the data well (e.g., **Fig. 3**), it is also important to note that several alternative versions of the DD or ID models did not fit the data (**Suppl. Tab. S2**). For example, the model assuming that BCG vaccination only reduces the rate of Mtb replication (*ϵ*_*r*_ *>* 0 and *ϵ*_*m*_ = 0) poorly fit the data (**Suppl. Fig. S6** and **Suppl. Tab. S2**); this important result suggests that reduced bilateral infection of the lungs of BCG-vaccinated mice is not a simple consequence of reduced CFU numbers (arising due to slower Mtb replication rate) and that BCG vaccination somehow impairs the ability of Mtb to disseminate in the lung (or outside of the lung). The model assuming that BCG vaccination increases Mtb death rate (with *ϵ*_*δ*_, **eqns. (1)–(2)**) also did not fit the data well (**Suppl. Tab. S2**). This result is somewhat consistent with our previous observation that in CD infection, the rate of Mtb replication changes with time since infection more often than death rate does^43^. Interestingly, however, allowing the the effect of BCG vaccination on Mtb death rate to be time-dependent (*ϵ*_*δ*_ = *ϵ*_*δ*_(*t*)) and allowing for the vaccine to reduce the Mtb dissemination rate (with *ϵ*_*m*_) did result in substantially improved the model fit (**Suppl. Fig. S7**C&D, AIC = 234.06). This model fit predicted that vaccine efficacy at increasing Mtb death rate declines over time reaching highest values early in infection; the model also predicted a large reduction in Mtb dissemination rate (**Suppl. Fig. S7**C&D). Thus, this version of the model suggests that BCG vaccination may influence the rate at which Mtb is being eliminated during early stages of infection.

### Stochastic simulations of *L*_1_ → *L*_2_ version of the DD model fails to reproduce observed CFU variability

One of the key features of CFU data from ULD-infected mice is their large variability that typically is not observed in CD-infected mice^42^. When fitting ODE-based models to such variable data we make an implicit assumption that the data variability comes from measurement noise; however, given that in these experiments infection starts with very few bacteria (**Suppl. Fig. S1**A), variability in cell numbers may also arise due to stochasticity of Mtb replication, death, and migration (i.e., process noise)^42^. We therefore sought to investigate whether stochastic simulation of Mtb dynamics using best fit parameters of a model would match CFU variability measured experimentally.

To simulate Mtb dynamics stochastically we 1) converted the deterministic, ODE-based *L*_1_ → *L*_2_ version of the DD model into a set of transition rules (**Tab. 1**); 2) used parameters from the best fits of the DD model to the data from unvaccinated or BCG-vaccinated mice (**Fig. 2**A-B and **Fig. 3**A-B); 3) set the initial conditions as *L*_1_(0) = 1 and *L*_2_(0) = 0, and 4) ran 100 Gillespie simulations (and see Materials and methods for detail) for each of a set of model parameters for unvaccinated and BCG-vaccinated mice (**Fig. 4**). As expected, stochastic simulations predicted divergent trajectories for CFU for individual runs with the average predicted CFU somewhat matching experimentally observed values; however, it was clear that there were some data points that stochastic trajectories did not describe. In particular, while the model relatively well match variability in CFU numbers in the Lung 1, it failed to accurately predict larger CFU values in Lung 2 (**Fig. 4** B&D).

**Figure 4:**
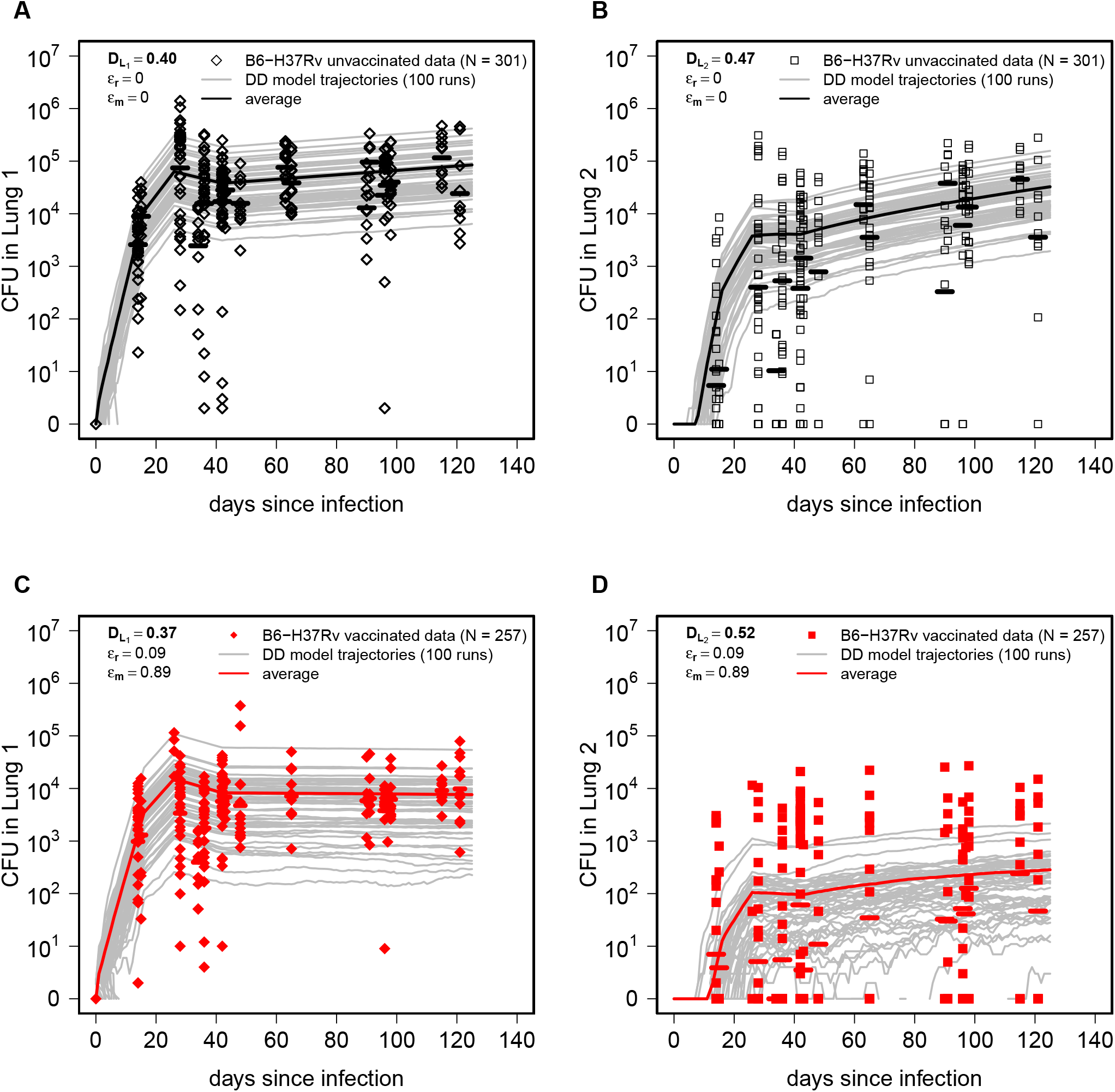
Stochastic simulations of the DD model fail to capture full variability of CFUs in the data. We translated the ODE-based DD model (**eqns. (1)–(2)**) into a corresponding set of transition rules (see Materials and methods for detail) and performed stochastic simulations using the best-fit parameter estimates (**Figs. 2 and 3**) for both unvaccinated (panels **A-B**) and BCG-vaccinated (panels **C-D**) mice. Gray lines denote individual runs of the model (total *n* = 100 runs) and solid lines denote the average of all simulated trajectories. To compare how well stochastic simulations match experimental data we calculated the Kolmogorov–Smirnov (KS) test statistic *D* (shown on individual panels) denoting the distance between cumulative distribution of CFUs predicted by the model and that observed in the data (**eqn. (9)**, see Materials and methods for detail).

To more rigorously evaluate the match between stochastic simulations and experimental data we used Kolmogorov-Smirnov (**KS**) test that measures distance between two distributions (**eqn. (9)** and see Materials and methods for detail). Similarly to the visual comparison, KS test metric was higher for model predictions in the Lung 2 as compared to Lung 1, and in all cases *D* values were relatively high (*D* = 0.37 − 0.52) suggesting that stochastic simulations of the *L*_1_ → *L*_2_ version of the DD model were not able to fully capture variability of experimental data.

### Realistic stochastic simulations of infection of right and left lungs better capture CFU variability

The mismatch between stochastic realizations of the DD model and experimental data may have arisen because of simplifying assumptions in the modeling framework. First, in the DD model we assumed that infection starts with a single bacterium; however, while this is probably correct for most ULD infections with an effective dose of 1 CFU/mouse, there is a substantial chance that two or three bacteria may initiate the infection^42^. Second, the DD model assumes that infection always starts in Lung 1; however, we have previously established that when the number of founding bacteria is greater than one, both lungs may be infected with the right murine lung having about 2 fold higher probability of infection (**Suppl. Fig. S1**B). The latter assumption is likely to result in under-predicted numbers of bacteria in the Lung 2 (**Fig. 4**B&D).

We therefore extended stochastic simulations of the DD model to include additional biological detail of ULD infection (dubbed as “realistic stochastic simulations”). First, we introduced random sampling of the actual number of infecting bacteria from the Poisson distribution assuming that the average dose is 1 CFU/mouse (**Suppl. Fig. S1**A). In case when the sampled number is above zero, we partitioned the sampled bacteria between right or left lung with 2/3 : 1/3 probability (**Fig. 1**C). We then modelled Mtb growth in the right or left lung with estimated replication (*r*_*L*_) and death (*δ*) rates, and possibility of migration between the lungs at rate *m* (**Figs. 2 and 3**) assuming direct dissemination between the lungs (**Fig. 1**D).

Importantly, the realistic simulations of Mtb dynamics provided a larger range of predictions of bacterial numbers for both right and left lungs and more accurately predicted both large and small CFU numbers in BCG-vaccinated mice (**Fig. 5**). The improved visual match of the data was confirmed by lower values of KS statistics (compare **Fig. 4** and **Fig. 5**). Yet, *D* values remained relatively high suggesting that this model still did not fully capture CFU variability observed experimentally. Another interesting observation was that the average CFU predicted by the model did not accurately match averages observed in experiments (**Fig. 5**) suggesting that parameter estimates generated in the DD model may not be fully accurate for the realistic stochastic simulations.

**Figure 5:**
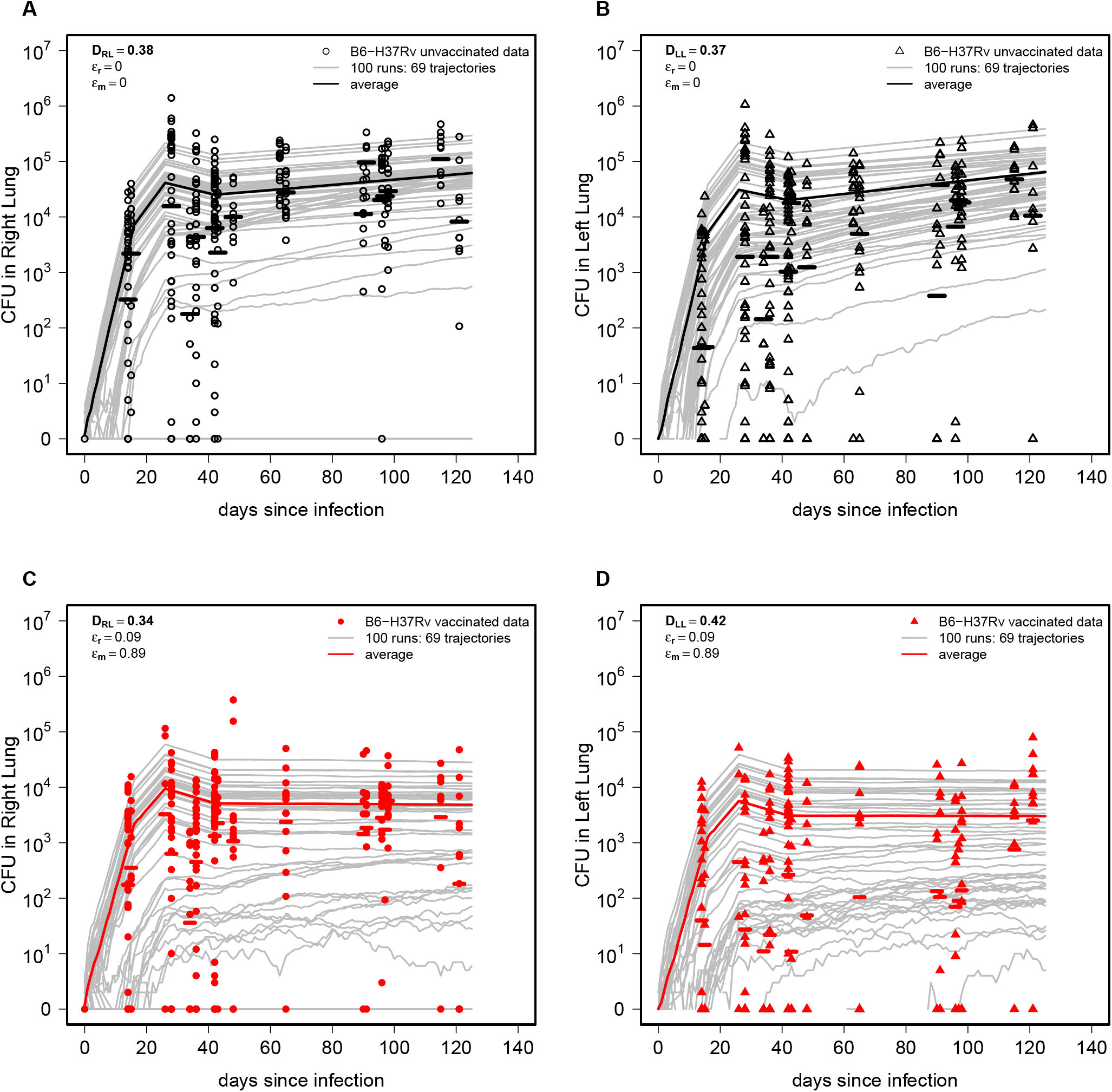
Stochastic simulations of Mtb infection, growth and dissemination between right and left lungs better match experimental data. Using the estimated parameters from the best fit DD model (**Figs. 2 and 3**), we ran the stochastic simulations assuming random infection of the right and left lungs and dissemination between the lungs (**Fig. 1**C-D) for unvaccinated (**A-B**) or BCG-vaccinated (**C-D**) mice. In simulations, we randomly chose the number of bacteria initiating infection from a Poisson distribution with an *λ* = 1 (**Suppl. Fig. S1**A), and used a probability of depositing a bacterium in the right lung of 66% (**Fig. 1**C and **Suppl. Fig. S1**B). We quantified the quality of the model match of experimental data using KS statistics *D* (**eqn. (9)** and Materials and methods for more detail).

### Hypothetical vaccines reducing Mtb replication rate more effectively limit Mtb dissemination

The DD model predicted that BCG-induced reduction in the rate of Mtb replication results in smaller lung CFU, and, as a consequence, in reduced frequency of bilateral lung infection; however, this was insufficient to fully explain smaller CFU in the Lung 2 (**Suppl. Fig. S6** and **Suppl. Tab. S2**). Having established that realistic stochastic simulations of the DD model relatively well explain variability of bacterial numbers in the lungs of unvaccinated or BCG-vaccinated mice we sought to investigate how a hypothetical vaccine that exclusively reduces the rate of Mtb replication or the rate of Mtb dissemination between the lungs, would influence the frequency of bilateral infection.

As expected increasing efficacy of a vaccine blocking the rate of Mtb replication *ϵ*_*r*_ reduced the frequency of bilateral infection (**Fig. 6**A); interestingly, the impact of replication rate-reducing vaccine was moderate for smaller vaccine efficacy (*ϵ*_*r*_ ≤ 0.3), and was relatively independent on the efficacy of the vaccine reducing the rate of Mtb dissemination *ϵ*_*m*_ (**Fig. 6**A). In contrast, increasing efficacy of a vaccine that reduces the rate of Mtb dissemination had relatively minor impact on the frequency of bilateral infection unless vaccine efficacy is relatively high (*ϵ*_*m*_ *>* 0.9, **Fig. 6**B). These results suggest that vaccines that reduce the rate of Mtb replication even slightly should have stronger impact on the ability of Mtb to disseminate between the lungs than vaccines that only reduce ability of Mtb to disseminate between the lungs.

**Figure 6:**
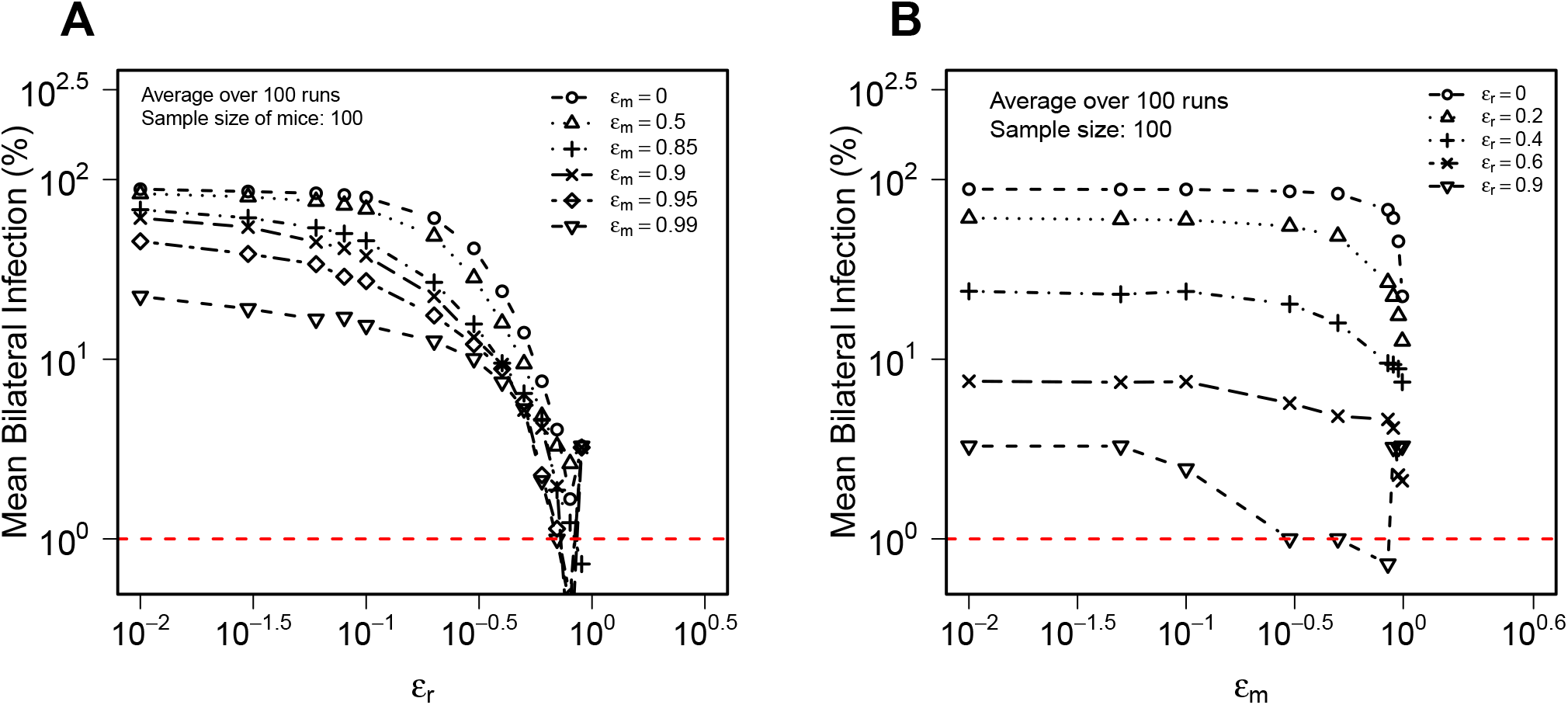
A vaccine reducing the rate of Mtb replication has a greater impact on probability of Mtb dissemination in the lungs as compared to a vaccine that exclusively blocks Mtb dissemination. We run stochastic simulations of Mtb replication in and dissemination between RL and LL (**Fig. 1**C-D) using parameter estimates of DD model fit to CFU data in unvaccinated mice (**Fig. 2**) for 100 mice vaccinated with a vaccine that reduces the rate of Mtb replication (*e*_*r*_) or a vaccine that reduces the rate of Mtb dissemination between lungs (*e*_*m*_); note that model predictions for *e*_*r*_ = *e*_*m*_ = 0 denote unvaccinated mice (value 10^−2^ on the *x* axis actually corresponds to *e*_*r*_ = 0 or *e*_*m*_ = 0). We calculate the percent of mice that have detectable CFU in both lungs (i.e., with bilateral infection) at day 14 post-infection with limit of detection LOD = 1 for varying levels of *e*_*r*_ (**A**) or *e*_*m*_ (**B**). We performed 100 runs per one set of parameters and show the average percent of bilateral infection.

### Power analysis for detecting the vaccine efficacy

To detect small efficacy of BCG vaccine at reducing probability of infection of mice exposed to ULD dose, Plumlee *ϵt al*. ^42^ had to combine data from 21 experiments, in total involving over 1,000 mice. It was not clear, however, whether these experiments were powered enough to actually detect a small efficacy of the vaccine (∼10%) at preventing infections. In addition, it was not previously possible to also determine how many mice would be required to detect efficacy of a hypothetical vaccine that only reduces the rate of Mtb dissemination. We therefore used realistic stochastic simulations of the DD model to perform such power analyses, i.e., to investigate the number of mice that would be required to use in an experiment to detect efficacy of a vaccine that reduces the rate of Mtb replication *ϵ*_*r*_ or reduces the rate of Mtb dissemination in the lung *ϵ*_*m*_; this is similar to an approach we have outlined recently^55–57^.

We ran the model simulations (with dose = 1) by assuming 1) different numbers of mice per unvaccinated or vaccinated group, 2) different values of the vaccine efficacy *ϵ*_*r*_ or *ϵ*_*m*_, and 3) by calculating the proportion of “experiments” (out of 1000 simulations) that result in statistically significant difference in proportion of infected mice (for *ϵ*_*r*_) or difference in proportion of mice with bilateral infection (for *ϵ*_*m*_, see Materials and methods for detail). As expected, increasing the number of mice per group resulted in larger power to detect a given level of vaccine efficacy and fewer mice were needed to detect larger efficacy levels (**Fig. 7** and **Tab. 3**). In particular, relatively small numbers of mice (*n*_mice_ = 20) are needed to detect highly efficient replication rate-reducing vaccine (*ϵ*_*r*_ ∼0.7, **Fig. 7**A), and even at moderate levels of vaccine efficacy (*ϵ*_*r*_ = 0.5), only 50 mice (per group) would be needed to detect reduction in the frequency of infected vaccinated mice (**Tab. 3**). In contrast, dissemination-blocking vaccines would require relatively large numbers of mice (*n*_mice_ = 100) to detect high vaccine efficacy levels (*ϵ*_*m*_ = 0.90, **Fig. 7**B). This is perhaps unsurprising given that Mtb dissemination may occur continuously during the infection, and preventing dissemination to the collateral lung when CFU are sufficient large would require high vaccine efficacy. In addition, even in ULD infection setting, many mice would be infected with two or more bacteria in both lungs reducing the ability to detect efficacy of a dissemination-blocking vaccine.

**Table 3:** Calculation of number of mice needed to detect the efficacy of a hypothetical vaccine. We compared the sample size estimates required to achieve 80% power for different levels of vaccine efficacy, controlling replication (*ϵ*_*r*_) or dissemination (*ϵ*_*m*_). The power analysis is implemented by using realistic stochastic simulation to predict the sample size per group, as described in the Materials and methods section. Model-predicted sample sizes are calculated when measuring total lung CFU or bilateral infection at day 14 post-infection assuming dose = 1 and LOD = 1; we also list the sample size predicted in Plumlee *et al*. ^39^. Note that the previously reported sample sizes are for detecting infection probability that does not exactly correspond to our simulations assuming replication rate-reducing vaccine because infection probability is a nonlinear function of vaccine efficacy *ϵ*_*r*_ . “-” denotes not available.

| Category | Efficacy Value | Minimum sample size per group (from experiment<sup>39</sup>) | Model predicted sample size per group |
| --- | --- | --- | --- |
| Vaccination efficacy on replication ($\epsilon_r$) | 0.2 | 259 | 629 |
|  | 0.3 | 112 | 215 |
|  | 0.4 | 66 | 97 |
|  | 0.5 | 40 | 50 |
|  | 0.6 | 28 | 29 |
|  | 0.7 | 21 | 21 |
|  | 0.8 | 16 | 17 |
|  | 0.9 | 12 | - |
| Vaccination efficacy on dissemination ($\epsilon_m$) | 0.85 | - | 155 |
|  | 0.90 | - | 97 |
|  | 0.95 | - | 50 |
|  | 0.99 | - | 26 |

**Figure 7:**
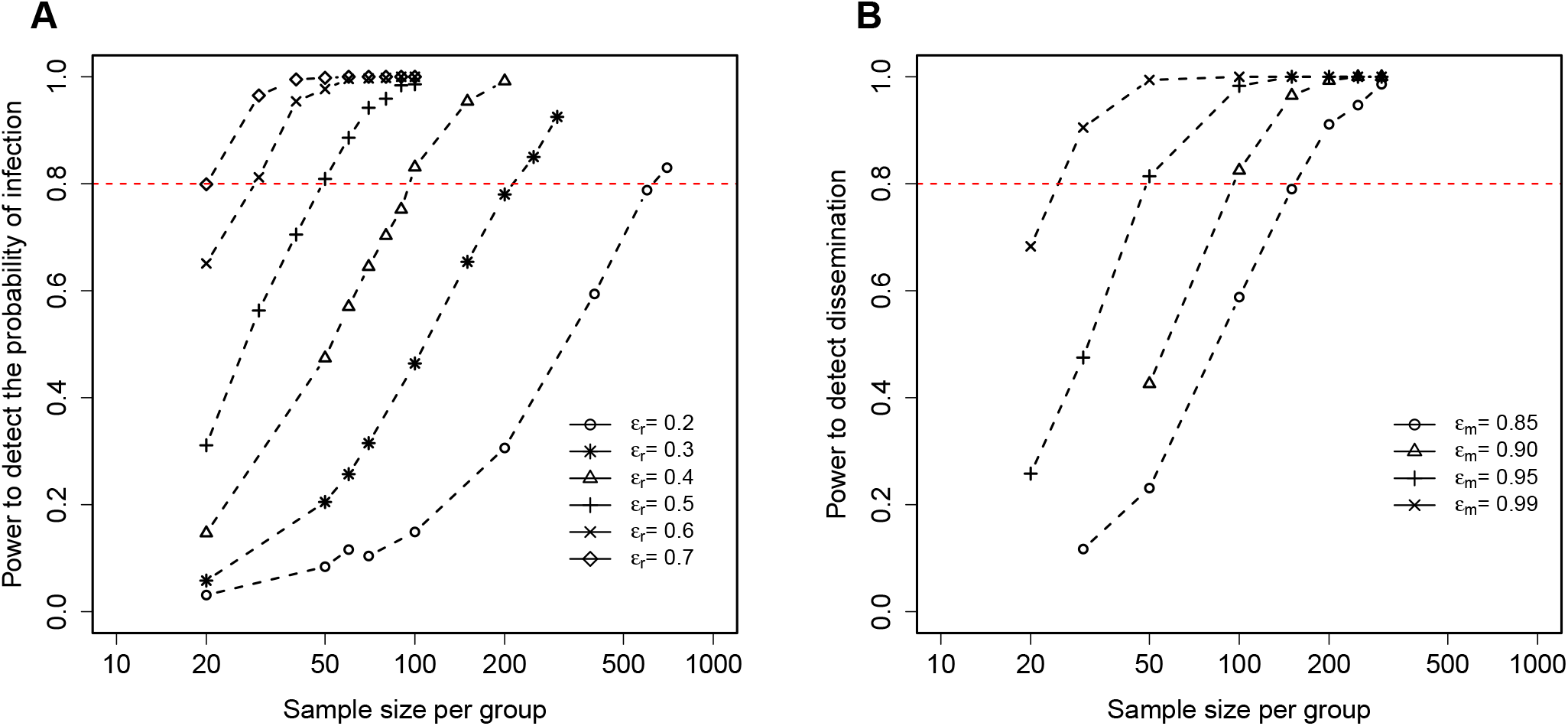
Realistic stochastic simulations of the DD model predict the number of animals required to detect efficacy of vaccines preventing Mtb infection or Mtb dissemination between lungs. We ran stochastic simulations of Mtb replication in and dissemination between RL and LL (**Fig. 1**C-D) in unvaccinated mice and mice vaccinated with a vaccine that either reduces the rate of Mtb replication (*ϵ*_*r*_, **A**) or reduces the rate of Mtb dissemination between the lungs (*ϵ*_*m*_, **B**) and calculated the proportion of mice that are uninfected (**A**) or have unilateral infection (**B**) by day 14 post infection assuming dose = 1 and LOD = 1. By varying the number of unvaccinated and vaccinated mice we calculated the probability of detecting vaccine efficacy of a particular value (*ϵ*_*r*_ and *ϵ*_*m*_ are noted on the panels, and see **Tab. 3**).

Plumlee *ϵt al*. ^39^ also performed power analysis predicting the minimal number of mice to be used in an experiment to detect efficacy of an infection-blocking vaccine in ULD-exposed mice. Interestingly, our mathematical modeling-based power analysis predicted somewhat larger numbers of mice required to detect efficacy of replication-blocking vaccine at preventing infection (**Tab. 3**). In particular, our simulations suggest that detecting a modest efficacy (*ϵ*_*r*_ = 0.2) would require 629 mice per group and well over 1,000 mice per group for *ϵ*_*r*_ = 0.1 (not shown). In additional analyses we found that measuring lung CFU at later time points (e.g., at day 35 p.i.) would require fewer mice to detect vaccine efficacy of a particular level – the required sample sizes decrease from 629 to 595 mice per group at *ϵ*_*r*_ = 0.2 for CFU measurements at day 14 vs. day 35, respectively. Because the previous study used CFU measurements from different time points while our power analysis was for a single time point, this could in part explain the difference in mouse number estimates. These results also suggest that later time points may provide greater statistical power to detect modest efficacy of infection-blocking vaccines. Furthermore, since we were able to detect a relatively small replication-reducing efficacy of BCG vaccine (*ϵ*_*r*_ = 0.09, **Fig. 3**) with ∼500 mice per group, this result indicates that CFU measurements at multiple time points may contain more information about vaccine-mediated inhibition of Mtb replication than a simpler binary infection outcomes. This prediction remains to be tested experimentally.

## Discussion

In this study, we developed and analyzed deterministic and stochastic mathematical models to investigate within-host Mtb dynamics at ULD infections of mice. By fitting direct and indirect dissemination models to experimental CFU data from Plumlee *ϵt al*. ^39^, we demonstrated that both models can successfully explain the observed Mtb kinetics, predicting rapid initial replication, transient control, and subsequent persistence (**Fig. 2**). After incorporating the effect of BCG vaccination, both models consistently indicated that BCG markedly reduces Mtb dissemination rate between lungs by approximately *ϵ*_*m*_ ≈ 90% while only modestly lowering bacterial replication rates (*ϵ*_*r*_ ≈ 9%, **Fig. 3** and **Suppl. Tab. S2**). Interestingly, we found similar estimates of BCG vaccine-induced reduction of Mtb replication rate by fitting a model that incorporates immune response-mediated control of Mtb growth to data from ULD or CD-infected mice (**Suppl. Fig. S8**). Realistic stochastic simulations of the DD model that capture randomness of actual number of bacteria inhaled by individual mice and possibility of bidirectional migration between right and left lungs relatively accurately reflect the variability of experimental CFU data (**Fig. 5**). Realistic stochastic simulations showed that a hypothetical vaccine that reduces the rate of Mtb replication has a larger effect at preventing bilateral infection as compared to a vaccine that only reduces Mtb dissemination rate (**Fig. 6**). Finally, our power analysis provided quantitative estimates of sample sizes needed to detect efficacy of vaccines blocking Mtb replication or dissemination (or both), offering a framework for optimizing preclinical study designs (**Fig. 7** and **Tab. 3**).

Our modeling results align closely with recent animal studies and clinical evidence supporting BCG vaccine-mediated protection against TB. In particular, our results are consistent with the experimental findings of Plumlee *ϵt al*. ^39^ where by infecting mice with barcoded Mtb strain the authors found reduced bilateral lung infection (by nearly 80%) in BCG-vaccinated mice. While their study quantified dissemination efficacy based on the frequency of bilateral infections, our modeling framework captures the underlying kinetics of Mtb replication and migration. Our conclusion that BCG vaccine-induced reduction in Mtb replication rate may be declining with time since infection (**Suppl. Fig. S7**) is consistent with observation in humans documenting decline in BCG vaccine efficacy over time^54^, although the kinetics of decline in vaccine efficacy does differ between mice and humans. Collectively, previous experimental studies and our results converge on a common interpretation: BCG provides partial but consistent protection that limits bacterial dissemination and disease severity rather than fully preventing the Mtb infection^58^.

Clinical and epidemiological data also support this interpretation. Decades of human studies, from early randomized placebo-controlled trials in the UK and Scandinavia to more recent trials such as Chingleput in India, have revealed substantial variability in BCG efficacy ^9,11^. Despite this variability, meta-analyses consistently show that BCG offers strong and durable protection against severe and disseminated forms of TB in children, including miliary and meningeal TB, while offering variable protection against adult pulmonary disease^14,15^. Though the reason behind this variable efficacy of BCG remains elusive, our analysis reveals a key mechanistic insight of a hypothetical vaccine: control of Mtb replication strongly governs dissemination outcomes, suggesting that vaccines that suppress replication are likely to provide broader and more consistent protection against pulmonary spread than those targeting dissemination alone. Thus, our work provides a predictive and quantitative framework that bridges experimental and clinical observations, reinforcing the concept that the primary impact of BCG vaccination lies in its ability to limit systemic spread of Mtb.

Due to lack of longitudinal measurements of immune cells in ULD-infected mice in our experimental data we opted for a simplified mathematical modeling framework where impact of immunity of Mtb dynamics is reflected via time-dependent Mtb replication and death rates (**eqns. (3)–(4)**). The estimated impact of BCG vaccination on these rates via constant or time-dependent parameters *ϵ*_*r*_ or *ϵ*_*δ*_ suggest hypotheses of how the vaccine impacts within-host Mtb dynamics^59^.

A reduction in the Mtb replication rate due to BCG (*ϵ*_*r*_) may reflect direct intracellular growth restriction within BCG-primed or IFN*γ*-activated macrophages. It is important to note that we detect impact of BCG vaccination on Mtb growth very early after infection (first 10-14 days, **Suppl. Fig. S8**) prior to arrival of Mtb-specific T cell response suggesting that BCG vaccination influences lung innate immunity including alveolar macrophages, via epigenetic reprogramming^60^, orr through early, T cell-independent recruitment of monocytes and neutrophils^61^. However, some these mechanisms remain actively debated in the field, and our model, by design, does not attempt to distinguish between them. Intracellular growth restriction might occur through phagosome acidification, nutrient or iron starvation, induction of phenotypic quiescence, or reduced Mtb burst size in macrophages^60,62^. It is also interesting to note that in the model with time-dependent *ϵ*_*r*_ the major impact of BCG vaccination is in the first 26 days of infection, with minimal contribution in the chronic phase (**Suppl. Fig. S7**), further highlighting the role of innate immunity in Mtb control.

Conversely, an increased *ϵ*_*δ*_ due to BCG vaccination may indicate heightened bactericidal capacity (via reactive oxygen/nitrogen species or phagosome-lysosome fusion)^63^ or CD8^+^ T-cell-mediated cytotoxicity; though the latter’s contribution may be constrained by the limited capacity of CD8^+^ T cells to recognize Mtb-infected macrophages in vivo^60,64^. Notably, the models with constant *ϵ*_*δ*_ fit the data much worse compared to those with BCG impact on replication (**Tab. S2**) making the outlined biological mechanisms less likely.

Finally, a reduction in dissemination rate (*ϵ*_*m*_) due to BCG vaccination may reflect enhanced tissue-level physical and cellular barriers, including accelerated granuloma encapsulation and impaired trafficking of infected phagocytes to distal sites such as another lung^65,66^. Our recent work suggests that migration of Mtb-infected cells from the lung to lung-draining lymph nodes is not prevented by BCG vaccination^65^; therefore, our predicted reduction in Mtb dissemination between the lungs may come from ability of BCG-induced immunity to control/eliminate Mtb at the disseminated sites. Taken together, as total lung CFU dynamics reflect net population outcomes, our model quantifies the functional impact of these three parameters without disentangling their underlying cellular drivers.

Our modeling framework is subject to several limitations that should be considered when interpreting the results. We reformulated the problem of Mtb dissemination in ULD-infected mice as initial infection (with 1 CFU) in Lung 1 and Mtb dissemination to Lung 2; this resulted in a simpler modeling framework (e.g., **eqns. (1)–(2)**) but required re-arranging of the data into Lung 1 (larger CFU) and Lung 2 (smaller CFU, **Fig. S3**). While this approach provided a consistent framework for fitting models, assuming different pathways of Mtb dissemination, to data, it is a simplifying assumption that may not accurately capture all infection scenarios. Indeed, stochastic simulations of the DD (*L*_1_ → *L*_2_) model did not well describe CFU variability observed experimentally (**Fig. 4**). It is re-assuring, though, that realistic stochastic simulations of the DD (RL += LL) model with parameters estimated assuming unidirectional dissemination matched better variable CFU measured experimentally (**Fig. 5**) suggesting that our parameter estimates are reasonable. The alternative approach to model Mtb dynamics at ULD doses deterministically is to consider, for example, DD (RL += LL) model with the ensemble of initial conditions of infection starting in the left, right, or both lungs, and weighting the model solutions proportionally to the frequency of a given initial condition. Work in this direction is ongoing and preliminary results suggest similar conclusions as those reached with a simplified approach (**Suppl. Fig. S9**).

In our analyses we assumed that Mtb starts growing exponentially from the time it lands in the lung. However, our unpublished experimental data on CD infection of mice suggest that there may be a delay of few hours to few days before the bacteria enter the phase of exponential growth. We would expect that including a delay in Mtb growth may increase the estimated rate of Mtb replication during first days after infection but because details of Mtb activation in ULD-infected mice are unknown, we opted for a model that ignores such a delay in start of exponential growth phase.

When assuming time-dependent Mtb replication and death rates we selected particular time points at which the rates may change (**eqns. (3)–(4)**); this was driven in part particular features in our data (i.e., decline in CFU between day 26-28 and day 40-50) that was not observed in previous studies due to difference in timing of measurements^43,45^. We found that changing the time intervals does impact quality of the model fit of the data but typically did not influence dramatically estimates of model parameters including impact of BCG on Mtb replication and dissemination rates (results not shown). However, future studies may need to explore whether allowing time interval boundaries to include as model parameters may allow for a better fit.

In our preliminary analyses, we found that it was not possible to determine changes in both replication and death rates when fitting ODE-based models to experimental data, so we assumed Mtb death rates estimated previously^43,45^. We chose this approach because we estimated that death rate changes only once during the infection^43^. However, an alternative approach could be to fix the rate of Mtb replication to values found previously^43^ and estimate Mtb death rate by fitting the models to data. We will explore this possibility in our future work.

In our analysis we excluded uninfected mice; such exclusion could potentially bias parameter estimates toward higher infection probabilities and obscure vaccine effects on infection establishment. However, it is important to emphasize that the detected reduction in the rate of Mtb replication due to BCG vaccination *ϵ*_*r*_ = 0.09 translates to 9% increase in infection clearance probability (*p*_0_ = *δ/*(*r*(1 − *ϵ*_*r*_)) ≈ (1 + *ϵ*_*r*_)*δ/r* when *ϵ*_*r*_ ≪ 1 and for dose = 1) as was observed experimentally^39^. The models developed in the paper do not include explicit changes in immune components such as macrophages, neutrophils, or antigen-specific T cell response. In part, this was driven by the lack of experimental data on dynamics of these cell population in ULD-exposed mice. Extending the models to consider more mechanistic details of Mtb control in ULD infection is a key area of our current research.

We found that both direct and indirect dissemination models fit the experimental data with somewhat similar quality (**Suppl. Tab. S2**); yet, the DD model was the simplest and thus was favored by statistical metrics such as AIC. The ID dissemination model predicted slightly lower CFU values in an intermediate tissue as was observed experimentally in the spleen (**Suppl. Fig. S4**), highlighting the need to further improve the ID model. Additional data on Mtb burdens in other tissues, such as lung-draining lymph nodes, spleen, and potentially blood, should be able to narrow down specific pathways of Mtb dissemination in the lung. Notably, recent work has shown that BCG vaccination reduces Mtb burden in lung-draining lymph nodes without significant altering of myeloid trafficking or T cell priming, suggesting that BCG may not uniformly block dissemination but instead limits bacterial expansion and niche establishment at extrapulmonary sites ^65^.

In fitting the models to data we rescaled all CFU values by one, so that any zero CFU data become similar to the limit of detection of 1 CFU. Future work will need to explore if alternative treatments of zero CFU data, e.g., by assuming low CFU values follow a distribution^67^, result in different conclusions. Finally, our stochastic simulations have shown great variability in predicted lung CFU and yet, the ODE models we fit to data primarily utilizing average CFU values. Comparing predictions of stochastic simulations to data using KS test provides quantitative metric to evaluate fit quality but it remains to be explored if KS test could be used to rigorously fit stochastic models to data^50^.

Our work opens avenues for future research. There have been limited methods of comparing predictions of stochastic mathematical models to data. Our approach, based on calculating the average KS statistics (**eqn. (9)**), could be extended to minimize *D* with respect to model parameters in standard optimization routines^50^. Our work also highlights that impact of a vaccine could be determined relatively early after infection (**Suppl. Fig. S8**) and that time course CFU data may be more powerful at detecting efficacy of replication-blocking vaccines as compared to measuring proportion of infected mice (**Tab. 3**). Whether these inferences are repeated in testing of other vaccines in ULD-infected mice remains to be determined.

A growing number of TB vaccine candidates require small animal models for vaccine testing. One such model, the ULD infection murine model, has already demonstrated ability to infer more protection-related parameters than has been done in CD-infected mice^39,42^. Our novel mathematical modeling framework by mimicking ULD infection of mice will likely help interpret future data obtained from TB vaccine testing using the ULD murine model. For example, by using CFU burden measured at a few time points, our model should be able to predict vaccine efficacy by comparing results of realistic stochastic simulations with the data. Combining mathematical models of Mtb dynamics at ULD infection with experimental data may be a powerful novel way to access efficacy of next generation TB vaccines.

## Abbreviations

CFU: colony forming unit
Mtb: *Mycobactϵrium tubϵrculosis*
TB: tuberculosis
MDR: multi-drug resistant
ODE: ordinary differential equation
SSA: stochastic simulation algorithm
ULD: ultra-low dose
DD: direct dissemination
ID: indirect dissemination
LOD: limit of detection
CI: Confidence interval

## Data and code sources

The R codes and reformatted data are available on github:

**https://github.com/dipanjan754/BCG/vaccination/efficacy/.**

## Ethics statement

No animal work has been performed.

## Author Contribution

VVG and DC conceived the overall concept of the study and developed alternative models. DC performed data analysis, fitting models to the data, and stochastic simulations in the ULD framework. VVG performed simulations to fit models to the data in the conventional dose framework. DC wrote the first draft of the paper and all authors read, edited, and agreed on the final version.

## Acknowledgment

We would like to thank Plumlee *ϵt al*. ^39^ for making their data available with the publication and members of Urdahl lab for discussions of ULD infection and TB vaccines. We especially thank Kevin Urdahl for his valuable feedback on the manuscript. We further thank all members of the Ganusov lab for their feedback on previous versions of the manuscript.

## Supplemental Information

### ID model assuming intermediate tissue is blood

In the ID model, if blood is treated as an intermediate tissue, Mtb neither replicates nor dies in this compartment. Instead, blood will serve solely as a transport medium, carrying Mtb from Lung 1 to Lung 2. Thus, the ID model equations will be modified as mentioned below:

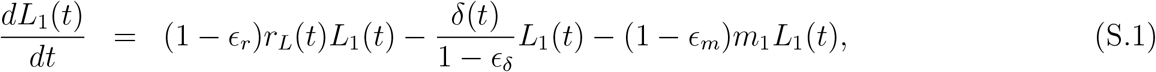

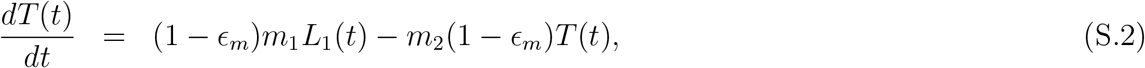

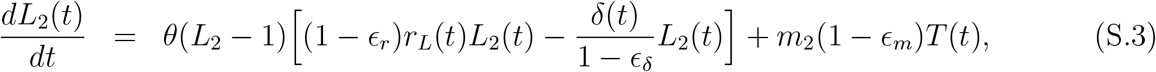

#### Modeling Mtb dynamics in the whole lung

To evaluate how BCG vaccination impacts early Mtb dynamics in mice infected with an ULD or conventional dose (**CD**) of Mtb, we modified our previously proposed mathematical model^65^ to include dynamics of immune response *ϵ* and suppression of Mtb replication by immunity^68^:

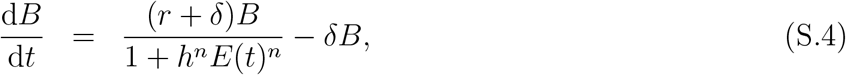

where *B* is the number of bacteria in the lung, *r* and *δ* are the rates of Mtb replication and death, respectively, *h* is the level of immunity at which immune-mediated suppression is half of its maximum, and *ϵ*(*t*) = e^*ρt*^ where *ρ* is the expansion of the immune response. We characterize kinetics of the immune response by time *T*_*ϵ*_ = − ln(*h*)*/ρ* that is the time when immunity reaches 50% of its suppressive capacity. We estimated *ρ* ≈ 0.4 − 0.6*/*day for Mtb-specific CD4 T cell response (Batabyal et al. (in prep)) but specific value of *ρ* is typically not very critical for the model behavior as long as it is sufficiently large^68^. In the model, infection starts with *B*_0_ bacteria.

### Calculation of effective dose in ULD-infected mice

Let *f*_0_ be the fraction of mice that remain uninfected, and *D* be the average dose of Mtb infection. Since Mtb infection is random, the number of bacteria inhaled by individual mice will follow a Poisson distribution^42,69^. Then, the probability that a mouse receives *k* bacteria is,

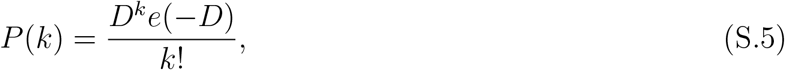

where the probability of no infection is: *P* (0) = *ϵ*^−*D*^ = *f*_0_. Thus, the effective dose *D* = − ln(*f*_0_). Note that the actual dose that mice are exposed to may be higher because of additional death rate of Mtb during the infection (**eqn. (3)**).

**Supplementary Figure S1:**
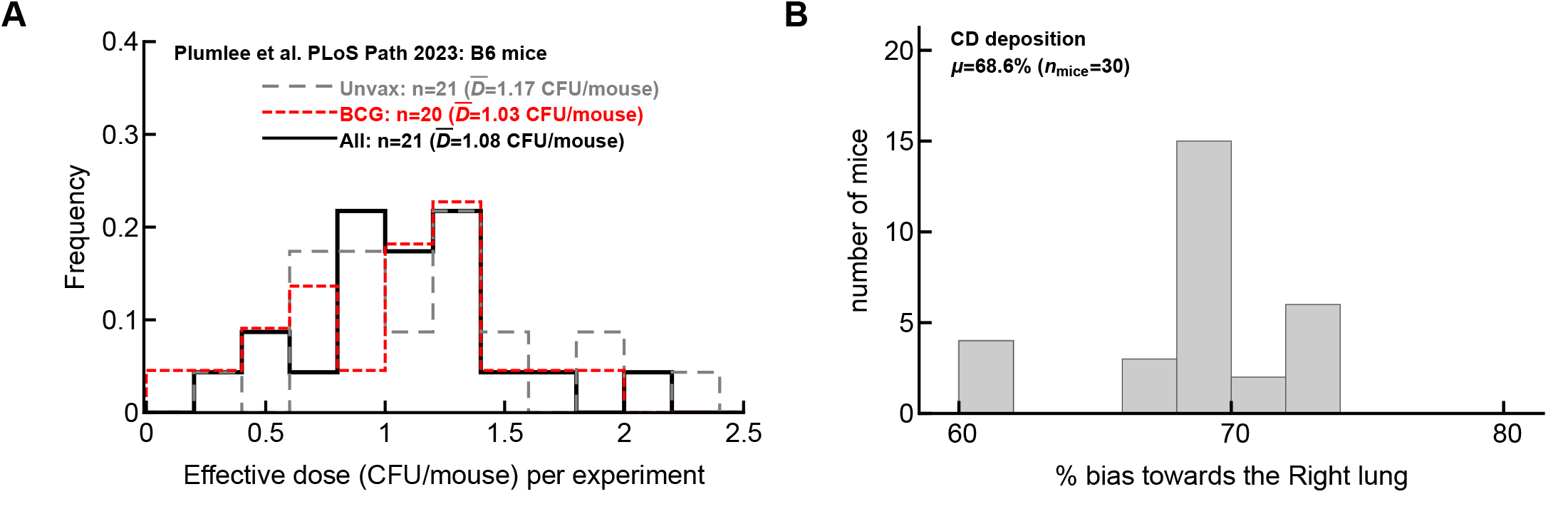
Distribution of estimated dose in mice exposed to ultra-low dose of Mtb and bias of initial infection towards the right lung of mice. **A**: For every experiment of *n* = 21 experiments of Plumlee *et al*. ^39^ we calculated the proportion of uninfected mice for unvaccinated group, BCG-vaccinated group, or all mice in the experiment (*f*_0_) and then calculated the average effective deposition dose using a Poisson distribution *D* = − ln(*f*_0_) (ref ^42^). In cases when all mice were infected, the dose could not be calculated, and such an experiment was excluded from the calculations. (Note a different number of experiments *n* used for calculation of the dose.) **B**: We calculated the bias in the CFUs recovered from right (CFU_RL_) or left (CFU_LL_) lung 1 day after Mtb infection of B6 mice with a conventional dose ^42^; the bias was calculated for each mouse as *b* = CFU_RL_*/*(CFU_RL_ + CFU_LL_).

**Supplementary Figure S2:**
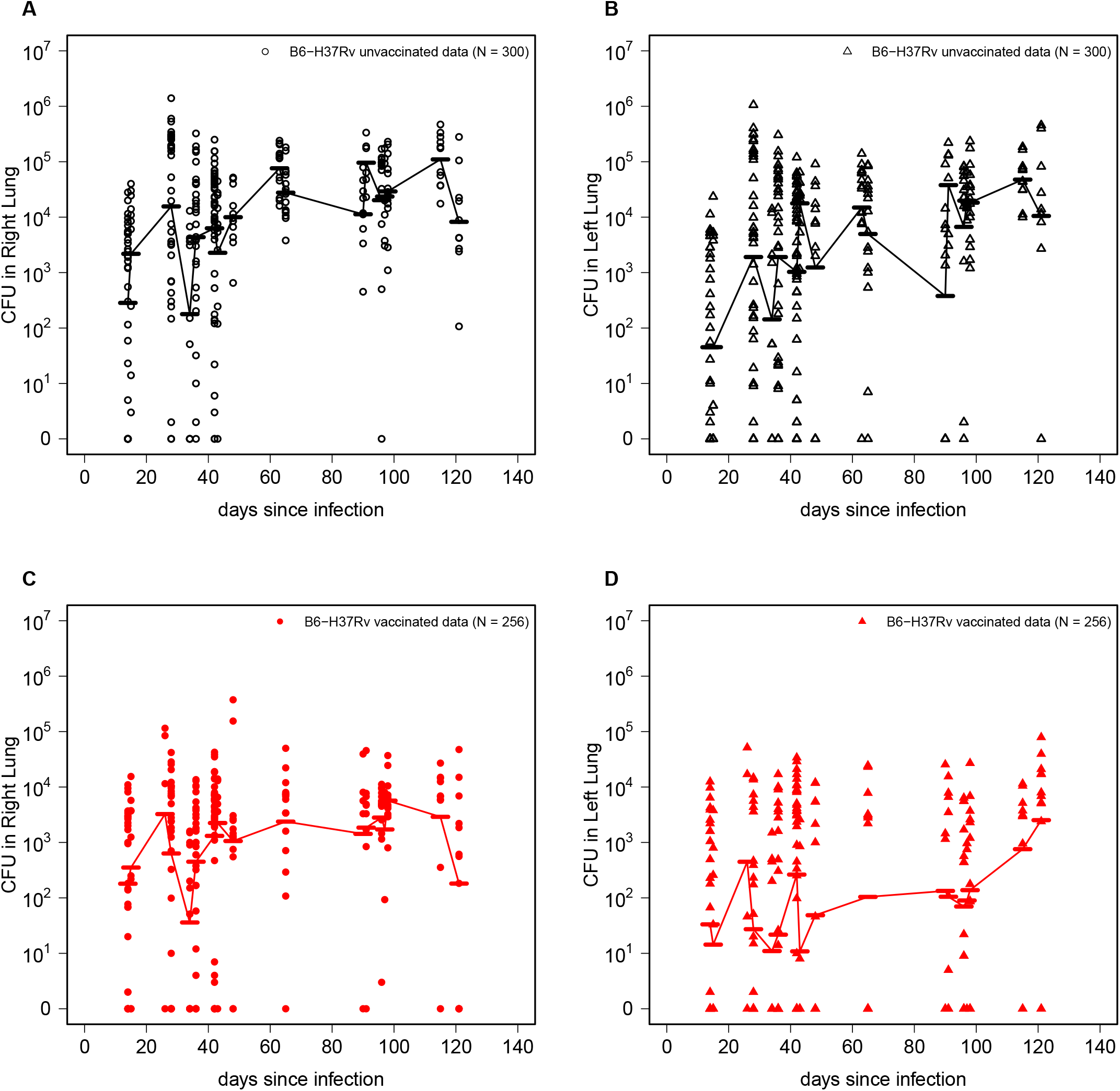
Bacterial burden at right lung and left lung of ultra-low dose infected mice Plumlee *et al*. ^**39**^. Mice were exposed to an ultra-low dose of Mtb H37Rv and bacterial burden at right lung (RL) and left lung (LL) was measured at different days post-infection after sacrificing the mice. Plumlee *et al*. ^39^ conducted multiple experiments, categorizing the mice into two cohorts: unvaccinated and BCG-vaccinated, comprising a total of (300 + 256) mice (excluding entries marked as ‘NA’ and the mice with no detectable infection i.e., mice with zero CFUs in both lungs or mice in which CFU was only measured in the whole lung without subdivision into right and left lungs). (A) and (B) present the CFU counts in the RL and LL of unvaccinated mice across all experiments, while (C) and (D) show the corresponding CFU data for BCG-vaccinated mice.

**Supplementary Figure S3:**
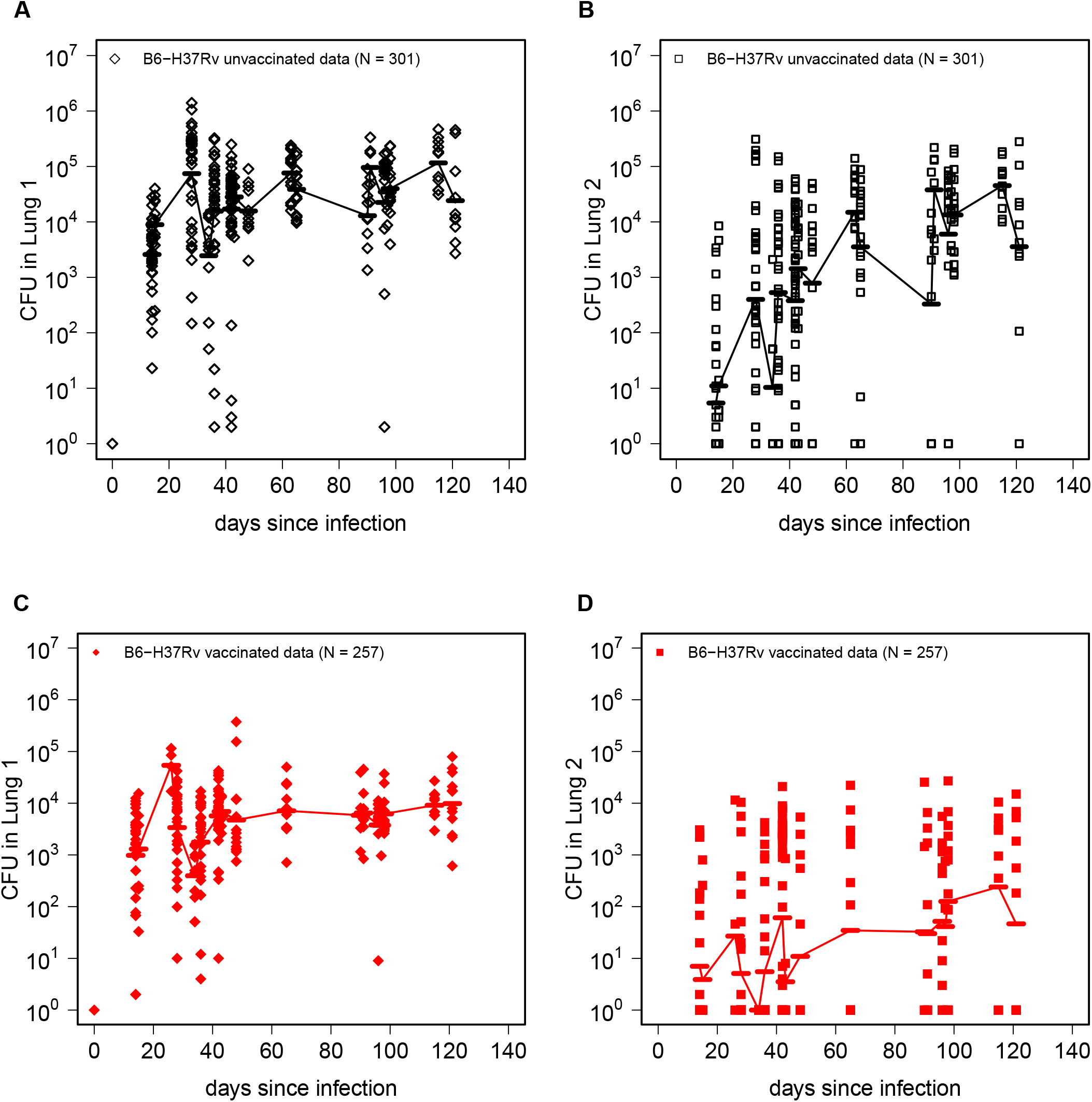
Reclassification of right lung and left Lung to Lung 1 and Lung 2. We have redefined the CFU data of right lung and left lung to Lung 1 and Lung 2 such that Lung 1 always has higher CFU than Lung 2. This reclassification reflects our assumption that the initial infection starts at Lung 1 (Lung 1 = 1 CFU at day 0), and then it could go to Lung 2 or other tissues as prescribed in the mathematical model section. Thus, due to the addition of an initial CFU data point, the number of mice in the dataset used for our analyses increases to 301 for unvaccinated mice (A, B) and to 257 for vaccinated mice (C, D).

**Supplementary Figure S4:**
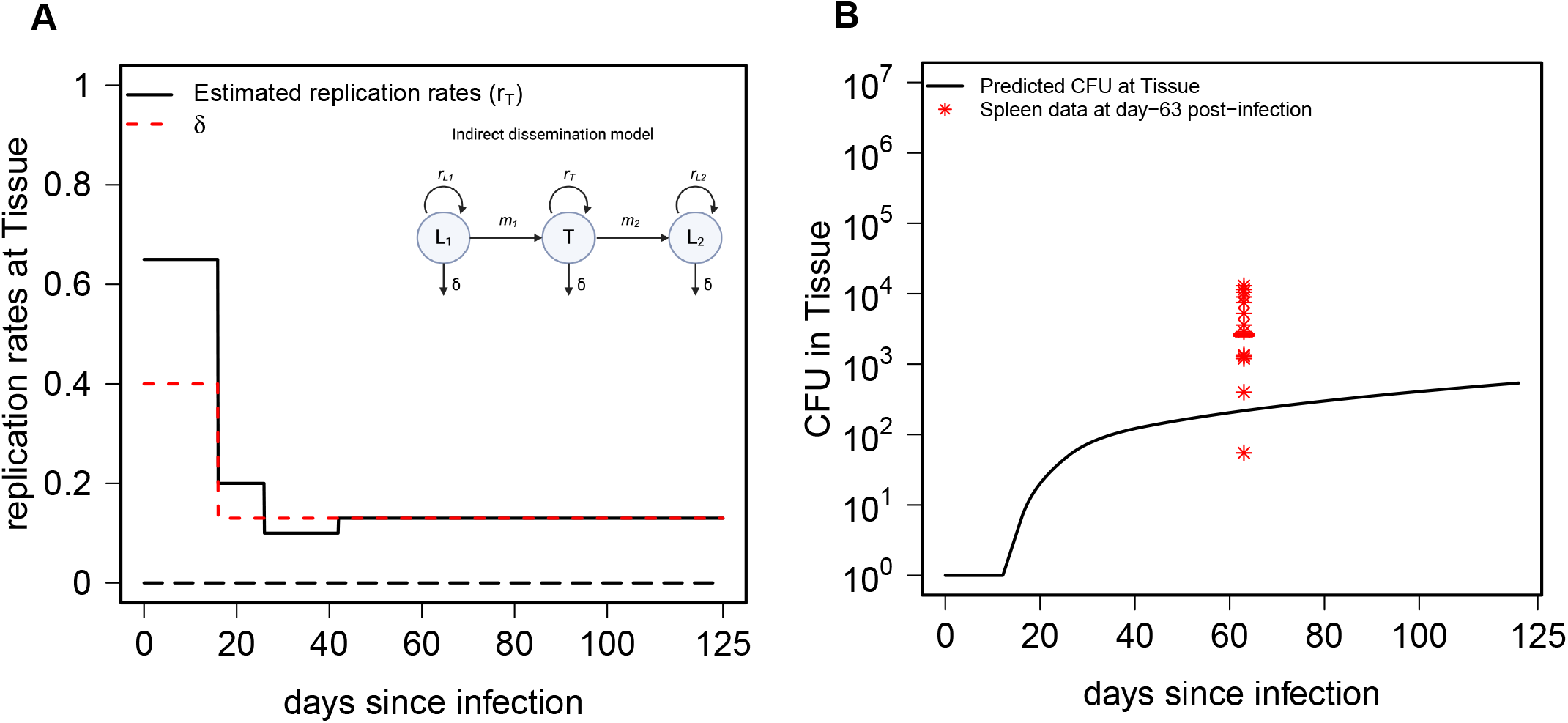
Estimated replication rates and predicted bacterial burden at an intermediate tissue predicted by the best fit of the ID model. **A**: We plot time-varying replication rates of Mtb in the intermediate tissue as estimated from fitting the indirect dissemination model to the data. **B**: We plot prediction of the model on the bacterial numbers in an intermediate tissue (line) and experimentally measured CFU in the spleen of ULD-infected mice (markers).

**Supplemental Table S1:**
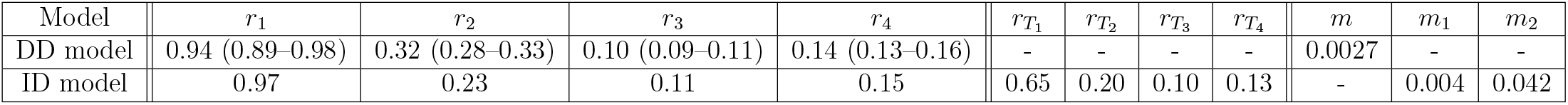
Estimates of replication rates of Mtb in the lung (*r*_*i*_) or lung (*r*_*i*_) and intermediate tissue 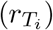, providing the best fit of the DD or ID model, respectively (model fits are shown in **Fig. 2**). The rates are time-dependent as defined in **eqn. (4)**. All rates are given in per day units, and − denotes not applicable. The 95% confidence intervals (CIs) for the parameter estimates were generated using modFit. We only report CIs for the DD model because modFit failed to provide CI estimates for the ID model due to the larger number of parameters.

**Supplementary Figure S5:**
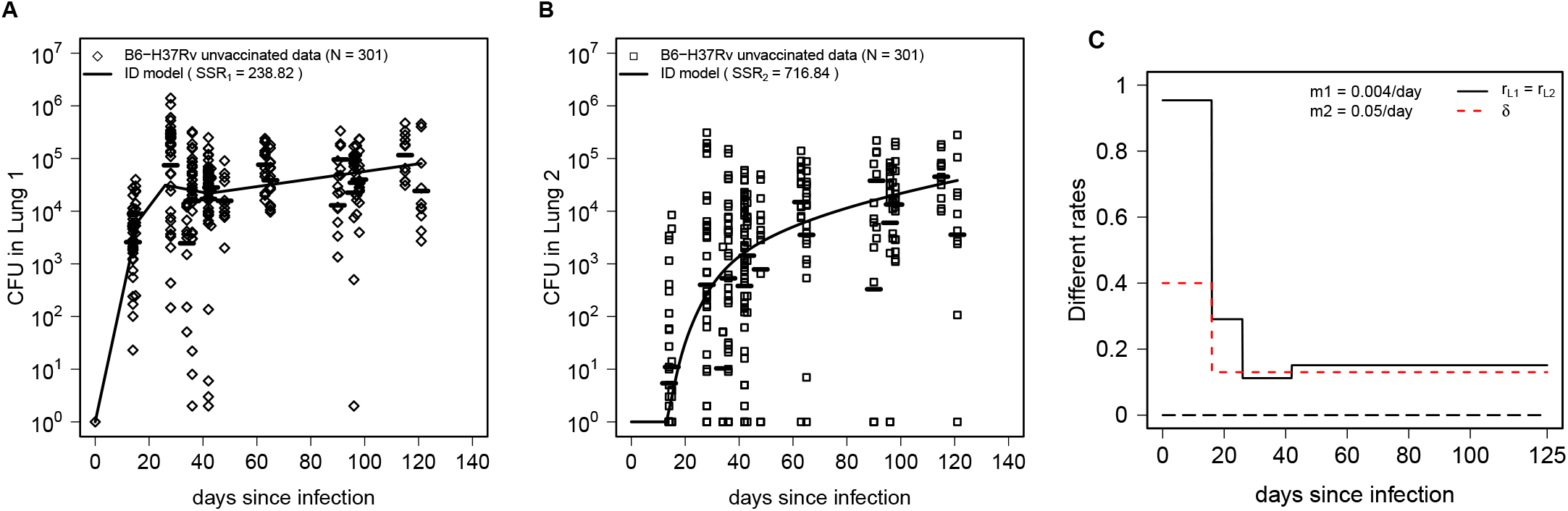
Indirect dissemination model explains the data well even if the intermediate tissue is assumed to be blood. We assume that in the ID model the intermediate tissue is blood, and there is no replication and death in the this intermediate tissue. Then, we fit the model to the data and the best-fitted model predicts the CFU burden at Lung 1 and Lung 2 (A, B). The best-fitted model estimate of the dissemination rate from blood to Lung 2 was *m* = 0.05*/*day, and the time-dependent Mtb replication rate in the lung behaves similarly to that in **Fig. 2**C.

**Supplementary Figure S6:**
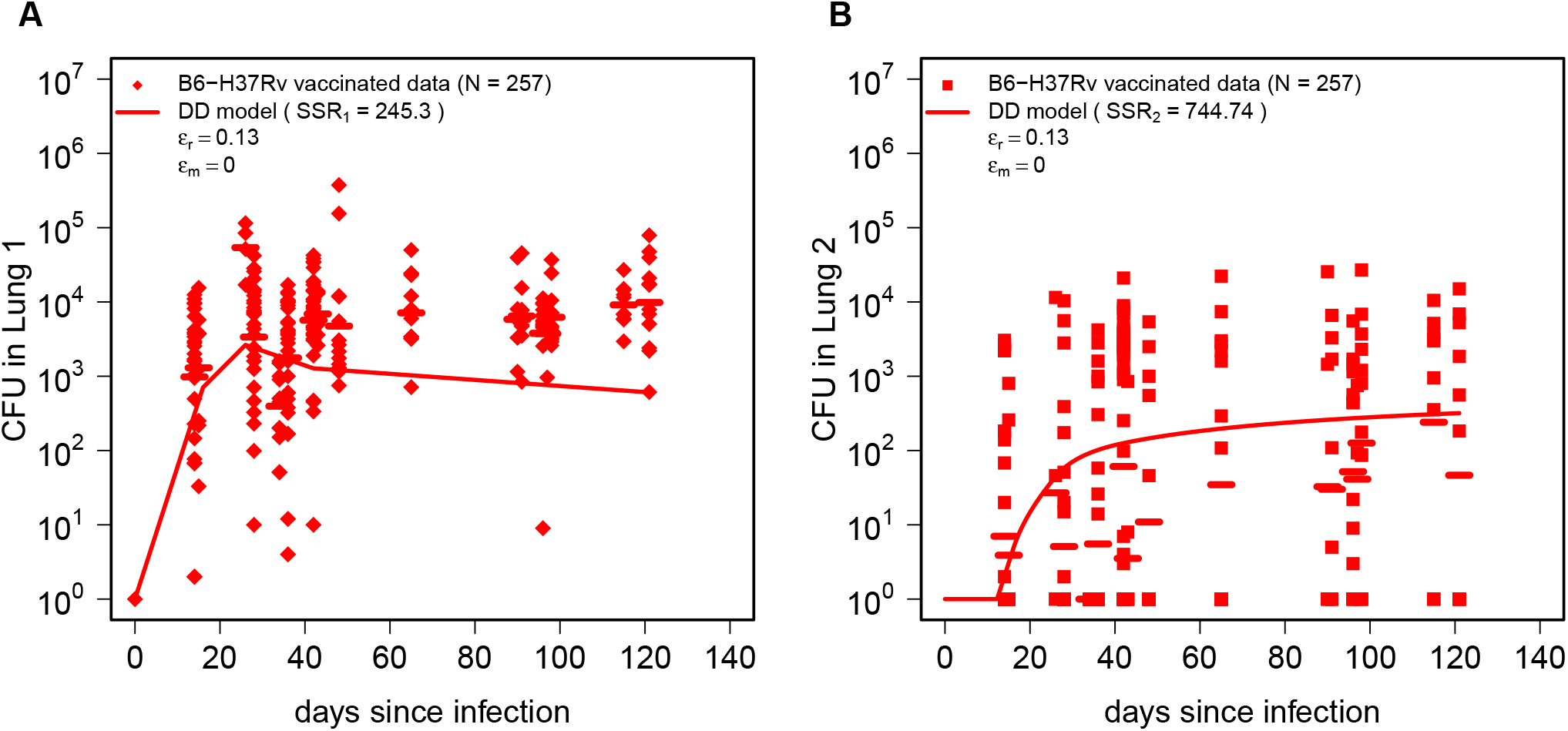
The DD model assuming that BCG only reduces the rate of Mtb replication does not accurately fit the data. We fit the DD model (**eqns. (1)–(2)**) to the BCG-vaccinated data by assuming that BCG vaccination only impacts the Mtb replication rate (*ϵ*_*r*_) but not the Mtb death rate and dissemination rate (*ϵ*_*δ*_ = *ϵ*_*m*_ = 0); estimated *ϵ*_*r*_ = 0.13. Other notations are similar to those in **Fig. 3**A-B. SSR values of the fit are shown on individual panels and AIC = 338.95 (see also **Suppl. Tab. S2**).

**Supplementary Figure S7:**
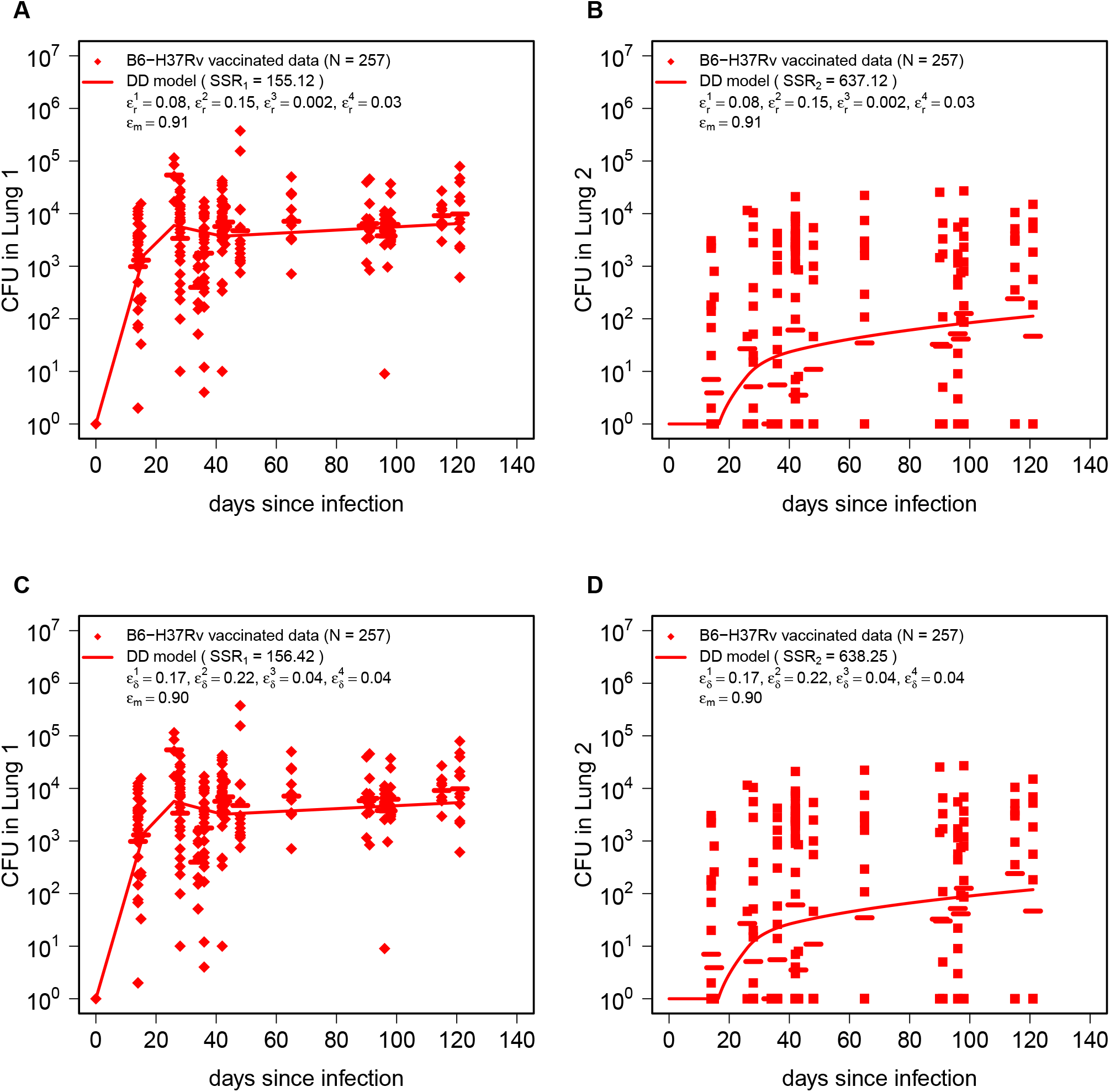
Efficacy of BCG vaccine declines with time since infection. We fit the DD model (**eqns. (1)–(2)**) to the data from B^1^ CG-vaccinated mice by assuming that BCG vaccine efficacy in reducing the rate of Mtb replication *ϵ*_*r*_ depends on time since infection in time intervals defined in **eqn. (3)**). The fit had much lower SSR (see **Suppl. Tab. S2**) and slightly lower AIC = 232.5 (A& B). Similarly, the DD model (**eqns. (1)–(2)**) is fitted to data from BCG-vaccinated mice, with the vaccine effect on the Mtb death rate (*ϵ*_*δ*_) modeled as time-dependent across intervals defined in **eqn. (3)**. The fit yields AIC = 234.06 (C & D).

**Supplementary Figure S8:**
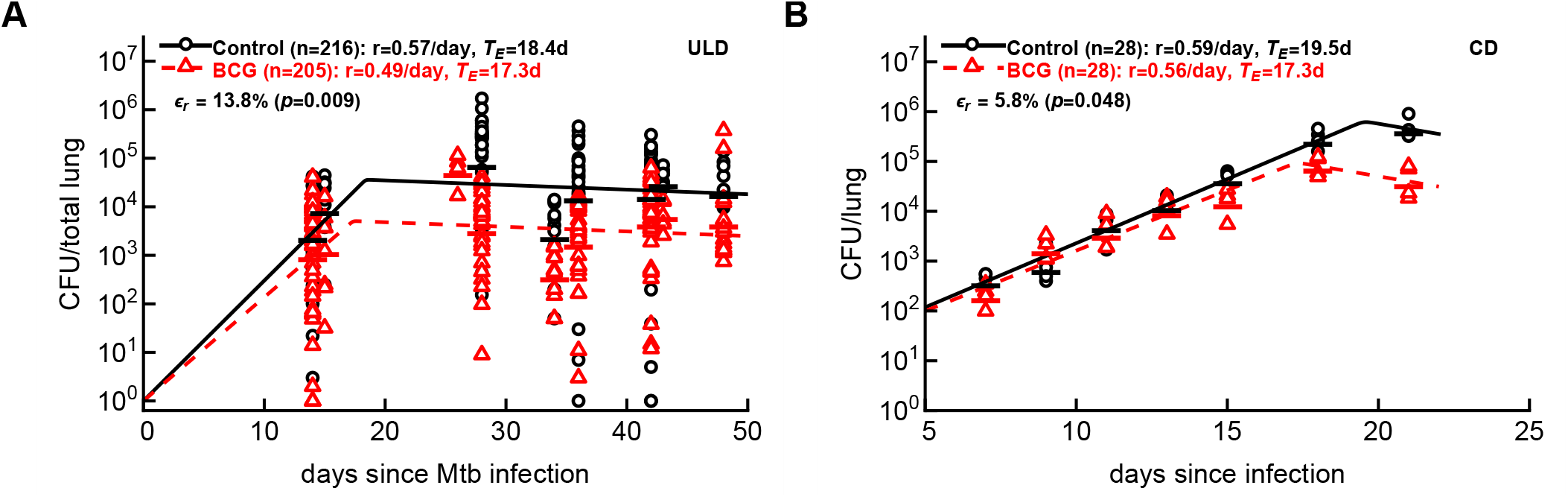
BCG vaccination reduces net Mtb replication rate early during ULD and CD infection of B6 mice. We fit our novel model that predicts Mtb dynamics in the presence of exponentially expanding immune response (**eqn. (S.4)**) to the data on Mtb numbers in lungs of B6 mice infected with an ultra-low dose (**A, Suppl. Fig. S2**) or conventional dose (**B**, ref ^70^) of H37Rv strain of mice. In panel A we only take the data the first 50 days of the total lung CFU of ULD-infected mice (**Suppl. Fig. S2**). We fit the model to data from both unvaccinated or BCG-vaccinated mice simultaneously only allowing for the parameters *r* and *h* (or *T*_*ϵ*_ = − ln(*h*)*/ρ*) to vary between the datasets; fixing the rate of Mtb replication between the datasets resulted in fits of poorer quality (A: *F*_1,416_ = 6.84, *p* = 0.009; B: *F*_1,49_ = 4.1, *p* = 0.048). Estimated efficacy of BCG vaccination was calculated as *ϵ*_*r*_ = (*r*_*unvax*_ − *r*_*BCG*_)*/r*_*unvax*_. Other estimated parameters are as follows. Panel A: *B*_0_ = 1 (assumed), *δ* = 0.22*/*day, *h*_*unvax*_ = 1.62 *×* 10^−5^, *h*_*BCG*_ = 3.02 *×* 10^−5^, *r*_*unvax*_ = 0.57*/*day, *r*_*BCG*_ = 0.49*/*day; panel B: *B*_0_ = 6.12, *δ* = 0.24*/*day, *h*_*unvax*_ = 8.22 *×* 10^−6^, *h*_*BCG*_ = 3.1 *×* 10^−5^, *r*_*unvax*_ = 0.59*/*day, *r*_*BCG*_ = 0.56*/*day. Parameters *ρ* = 0.6*/*day and *n* = 20 (**eqn. (S.4)**) were fixed in model fits to data.

**Supplementary Figure S9:**
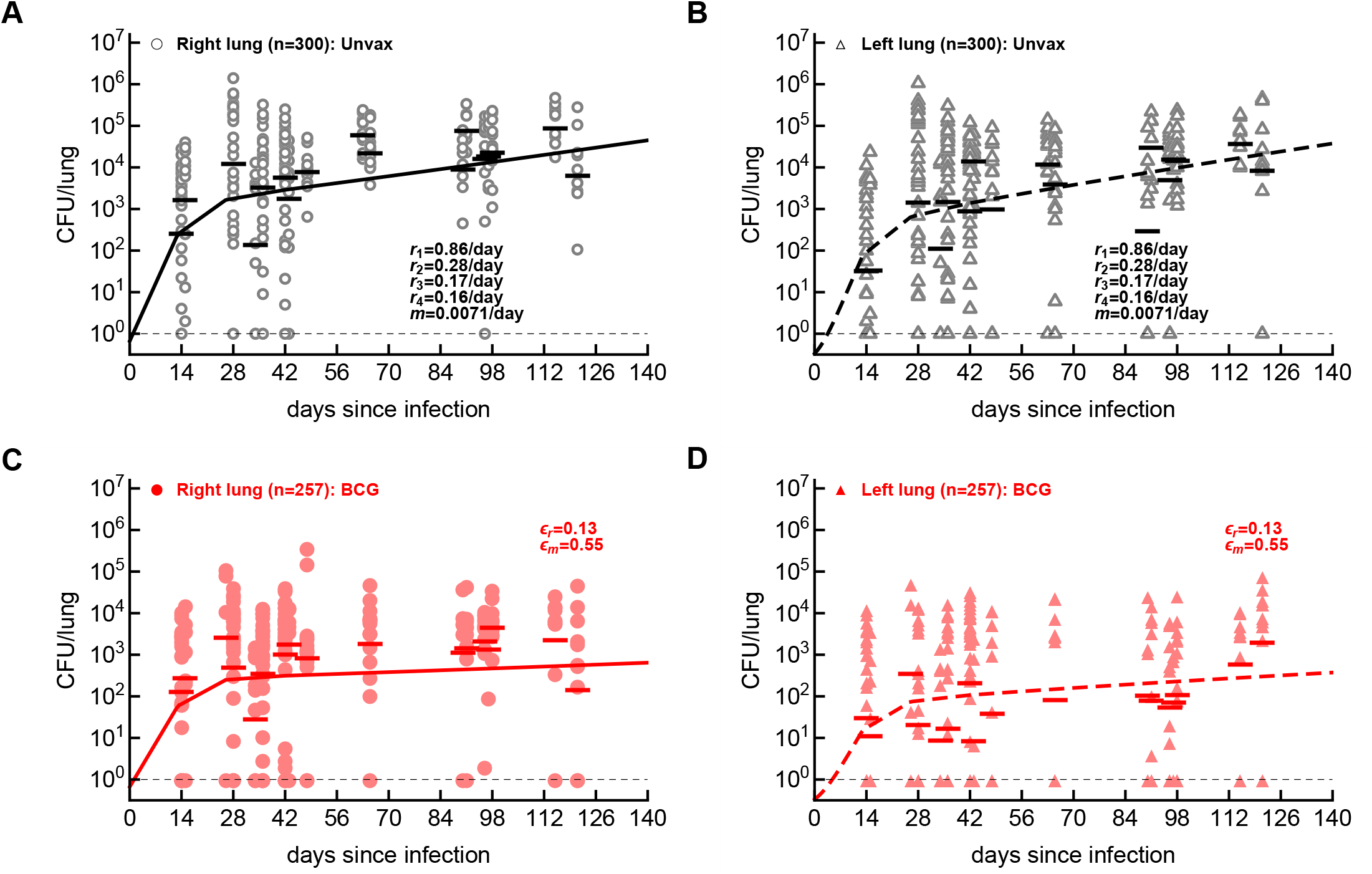
Deterministic version of realistic simulations of the DD model also predicts that BCG vaccination reduces Mtb replication and dissemination rates. We fit the DD model in which we consider random deposition of inhaled Mtb bacilli into right or left lungs, growth in the lungs, and dissemination between the lungs to the data from unvaccinated (panels **A-B**) or BCG-vaccinated (**C-D**) mice together assuming that BCG vaccine reduces the rate of Mtb replication (by *ϵ*_*r*_) and the rate of Mtb dissemination (by *ϵ*_*m*_). We plot experimental data (by markers) and model predictions (by lines); horizontal dashed line denotes the limit of detection of 1 CFU. Best fit parameters are shown on the panels and are *r*_1_ = 0.86*/*day, *r*_2_ = 0.28*/*day, *r*_3_ = 0.17*/*day, *r*_4_ = 0.16*/*day, *m* = 0.0071*/*day, *ϵ*_*r*_ = 0.13, and *ϵ*_*m*_ = 0.55; the death rates are assumed as in **eqn. (3)**. The best model fit resulted in SSR = 3494.14. To generate model predictions and fit the model to data, we solved the DD model (**eqns. (1)–(2)**) for a collection of initial conditions with an infection initiated by *n* bacteria (*n* = 1, 2 … *n*_max_) that distributed between right and left lung (e.g., (1, 0), (0, 1), (2, 0), (1, 1), (0, 2)… (*i, j*) for (RL, LL), respectively). For each of these initial conditions we calculated the probability *p*_(*i,j*)_ of such initial condition (*i, j*) happening using 0-truncated Poisson distribution and 2:1 bias in infection of the right:left lung. After solving the model for all initial conditions (e.g., assuming *n*_max_ = 3) we generated the average model predictions by weighting the solutions with a given initial conditions by their respective probabilities. To calculate the SSR, we use paired model predictions of the CFU number in the right and left lung and those observed experimentally; the residuals for a model prediction with initial condition (*i, j*) were squared and weighed by the probability *p*_(*i,j*)_ and summed over to generate SSR.

**Supplemental Table S2:**
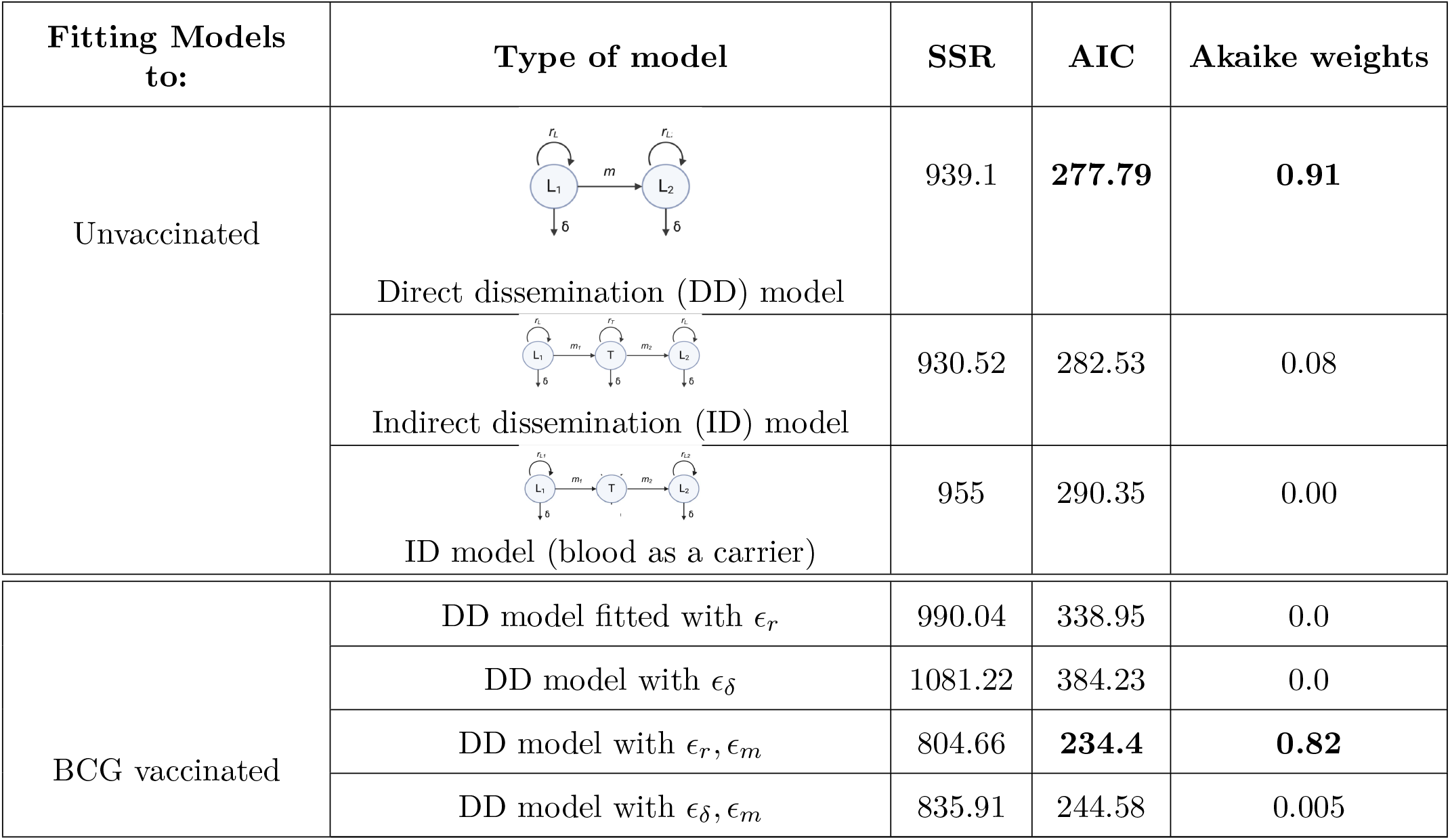
Comparison of model fits (SSR and AIC) for unvaccinated and BCG-vaccinated data. We show statistics of fits of alternative direct and indirect dissemination models, fitted to data from unvaccinated or BCG-vaccinated mice infected with ULD of H37Rv.

| Fitting Models to: | Type of model | SSR | AIC | Akaike weights |
| --- | --- | --- | --- | --- |
| Unvaccinated | Direct dissemination (DD) model | 939.1 | **277.79** | **0.91** |
|  | Indirect dissemination (ID) model | 930.52 | 282.53 | 0.08 |
|  | ID model (blood as a carrier) | 955 | 290.35 | 0.00 |
| BCG vaccinated | DD model fitted with $\epsilon_r$ | 990.04 | 338.95 | 0.0 |
| | DD model with $\epsilon_\delta$ | 1081.22 | 384.23 | 0.0 |
| | DD model with $\epsilon_r, \epsilon_m$ | 804.66 | **234.4** | **0.82** |
| | DD model with $\epsilon_\delta, \epsilon_m$ | 835.91 | 244.58 | 0.005 |

**Supplemental Table S3:** Calculation of number of mice needed to detect the efficacy of a hypothetical vaccine if the initial dose is *λ* = 2. We compared the sample size estimates required to achieve 80% power for different levels of vaccine efficacy, controlling replication (*ϵ*_*r*_) or dissemination (*ϵ*_*m*_). The power analysis is implemented by using realistic stochastic simulation to predict the sample size per group, as described in the Materials and methods section (and see **Tab. 3**).

| Category | Efficacy Value | Model predicted sample size per group |
| --- | --- | --- |
| Vaccination efficacy on replication ($\epsilon_r$) | 0.2 | 500 |
|  | 0.3 | 177 |
|  | 0.4 | 69 |
|  | 0.5 | 36 |
|  | 0.6 | <20 |
|  | 0.7 | <20 |
|  | 0.8 | <20 |
|  | 0.9 | <20 |
| Vaccination efficacy on dissemination ($\epsilon_m$) | 0.85 | 165 |
|  | 0.90 | 95 |
|  | 0.95 | 48 |
|  | 0.99 | 22 |

